# Biodegradation of the endocrine-disrupting compound bisphenol F by *Sphingobium yanoikuyae* DN12

**DOI:** 10.1101/2025.09.08.674859

**Authors:** Ruomu Chen, Yutian Gan, Wanting Huang, Chenyang Wang, Junhong Ge, Yuanyuan Cheng, Wenjing Qiao, Jiandong Jiang, Kai Chen

## Abstract

Bisphenol F (BPF) is an emerging environmental pollutant widely present in surface water and wastewater systems. Microbial activity is crucial in driving its degradation, offering a potential avenue for mitigating its environmental impact. Although the degradation pathway for BPF has been identified in various bacteria, the biodegradation mechanisms remain unclear. In this study, we isolated a highly efficient BPF-degrading strain of *Sphingobium yanoikuyae* DN12, which could utilize BPF as the sole carbon source and energy source for growth, from a river sediment in Anhui Province China. Through Ultra performance liquid chromatography high-resolution mass spectrometry (UPLC-HRMS) analysis, we found that oxidation and hydrolysis are key steps for BPF biodegradation. Utilizing whole-genome sequencing, comparative transcriptomics analysis and biochemical identification, a gene cluster *bpf* was identified to be involved in BPF degradation. BpfAB is a two-component oxidoreductase responsible for converting BPF to 4,4’-dihydroxybenzophenone (DHBP). BpfC is a Baeyer-Villiger monooxygenase (BVMO) responsible for converting DHBP to 4-hydroxyphenyl-4-hydroxybenzoate (HPHB). Isotope tracing demonstrated that the oxygen atom incorporated by BpfAB originates from water, whereas that incorporated by BpfC derives from molecular oxygen (O_2_). BpfD is an α*/*β hydrolase responsible for converting HPHB to 4-hydroxybenzoate (4HB) and 1,4-hydroquinone (HQ). Analysis of the taxonomic and habitat of 325 prokaryotic genomes revealed that BpfA-like homologs are predominantly found in the phylum *Pseudomonadota*, primarily inhabiting soil and aquatic environments. This study enhances our understanding of the biodegradation mechanism of BPF, and provides guidance for the effective remediation of BPF-contaminated environments.

**IMPORTANCE:** BPF is a widely used alternative to bisphenol A and poses a growing threat to ecosystems and human health due to its environmental persistence and endocrine-disrupting effects. Although microbial degradation pathways for BPF have been reported, the key enzymes involved and their catalytic mechanisms remain unclear. This work reports the isolation of a *Sphingobium* strain capable of mineralizing BPF and the genetic basis for the catabolic pathway. Three enzymes—a two-component oxidoreductase, a Baeyer-Villiger monooxygenase, and an α*/*β hydrolase—were biochemically characterized and shown to catalyze the three critical steps in BPF degradation. These findings provide insights into the biochemical processes involved in the microbial degradation of BPF.

## INTRODUCTION

Bisphenols (BPs), a class of endocrine-disrupting compounds (EDCs), have emerged as pollutants of growing concern in aquatic environments (1, 2). Extensive research has linked exposure to bisphenol A (BPA) with various health issues, including hypertension, obesity, type II diabetes, cardiovascular diseases, and cancer (3–5). As global regulations surrounding BPA use become stricter, the utilization of its substitute, bisphenol F (BPF), is increasing (6, 7). In 2018, global production of BPF reached hundreds of thousands of tons, primarily concentrated in developed regions such as North America, Europe, and Asia (8). BPF is a crucial raw material in the chemical industry and is widely used in the production of food can linings, coatings, adhesives, and electronics. However, BPF exposure has been associated with a range of toxicological effects, including developmental risks in children and reproductive toxicity in aquatic organisms (9–13). BPF residues are now commonly detected in environmental matrices such as industrial and municipal wastewater and landfill leachate (14–16).

Bioremediation, which primarily employs microorganisms, plants, or enzymes, is increasingly recognized as an effective method for degrading or transforming environmental pollutants (17). In recent years, an expanding number of BPF-degrading bacteria have been isolated, including *Pseudomonas* (18), *Sphingobium* (19–21), and *Bacillus* species (22), among others. A classic BPF biodegradation pathway has been identified in *Pseudomonas* and *Sphingobium*, initiating with the hydroxylation of the bridging carbon atom in BPF to form bis(4-hydroxyphenyl)methanol. This intermediate is subsequently oxidized to 4,4’-dihydroxybenzophenone (DHBP). An oxygen atom is then inserted between the keto carbon of DHBP and one of its benzene rings, converting the compound to 4-hydroxyphenyl-4-hydroxybenzoate (HPHB), and hydrolysis yielding 4-hydroxybenzoate (4HB) and 1,4-hydroquinone (HQ), ultimately leading to complete mineralization. Despite these advances, the degradation genes for BPF have not been identified, and molecular-level evidence of biodegradation is still lacking. Thus, elucidating the mechanisms of degradative enzymes underlying the degradation of BPF will enhance our understanding of BPF degradation pathways and broaden the repertoire of key enzymes capable of catalyzing BPs.

In this study, we utilized comparative transcriptomics, isotope labeling, and biochemical identification to reveal the key roles of a two-component oxidoreductase (BpfAB), a Baeyer-Villiger monooxygenase (BVMO) (BpfC), and a hydrolase (BpfD) in the degradation of BPF by strain DN12. These findings fill the gap in the research on the molecular mechanisms of BPF degradation by microorganisms, facilitate the prediction and assessment of the toxicity of intermediate and final products generated during BPF degradation, and provide important theoretical guidance and scientific evidence for designing efficient, safe, and sustainable remediation technologies for BPF pollution.

## RESULTS

### Isolation and identification of a BPF-degrading strain

A BPF-degrading bacterium, strain DN12, was isolated from the river sediment using a conventional enrichment culture technique. Colonies of strain DN12 on Luria-Bertani (LB) plates were smooth, creamy, oval, and Gram stain negative, with dimensions of 1.7 to 2.0 by 0.5 to 0.7 μm (Fig. S1A and S1B). Antibiotic susceptibility testing revealed that strain DN12 is resistant to streptomycin (Table S1). The phylogenetic analysis showed that strain DN12 was closely related to the *Sphingobium* species lineage and clustered with *Sphingobium yanoikuyae* ATCC 51230^T^ and *Sphingobium scionense* WP01^T^ (Fig. S1C), with sequence identity scores of 100% and 99.1%, respectively. The whole genome analysis of strain DN12 revealed four replicons, including one circular chromosome and three circular plasmids, pDN-1, pDN-2, and pDN-3 (Fig. S2; Table S2). The whole genome had an average G+C content of 63.2% and 5,176 predicted protein-coding sequences. The digital DNA-DNA hybridization (dDDH) between the genomes of strain DN12 (PRJNA1181918) and *Sphingobium yanoikuyae* ATCC 51230^T^ (PRJNA52201) was 72.6%, which is higher than the standard species boundary for dDDH (70%). Based on these results, the strain DN12 was identified as the same species as *Sphingobium yanoikuyae* ATCC 51230^T^, and named *Sphingobium yanoikuyae* DN12.

### Degradation of BPF by strain DN12

In minimal salts medium (MSM), strain DN12 was able to degrade 0.2 mM BPF to undetectable levels within 6 h, and cell growth was observed during the degradation process (Fig. 1A), indicating that strain DN12 can utilize BPF as the sole carbon and energy source for growth. Other BPs, such as BPA, bis(4-hydroxyphenyl)sulfone (BPS), tetrabromobisphenol A (TBBPA), and tetrabromobisphenol S (TBBPS), 2,2-bis(4-hydroxyphenyl)butane (BPB), bis(4-hydroxyphenyl)ethane (BPE), and bis(4-hydroxyphenyl)sulfide (TDP), were not degraded by strain DN12 (Table S3). The substrate-induced degradation experiment demonstrated that BPF-induced cells of strain DN12 degraded BPF at a significantly higher rate than uninduced cells (Fig. 1B), suggesting that the enzymes involved in BPF degradation are induced by BPF. Thus, transcriptome analysis can be an effective strategy to further explore genes related to BPF degradation.

**Fig. 1.**
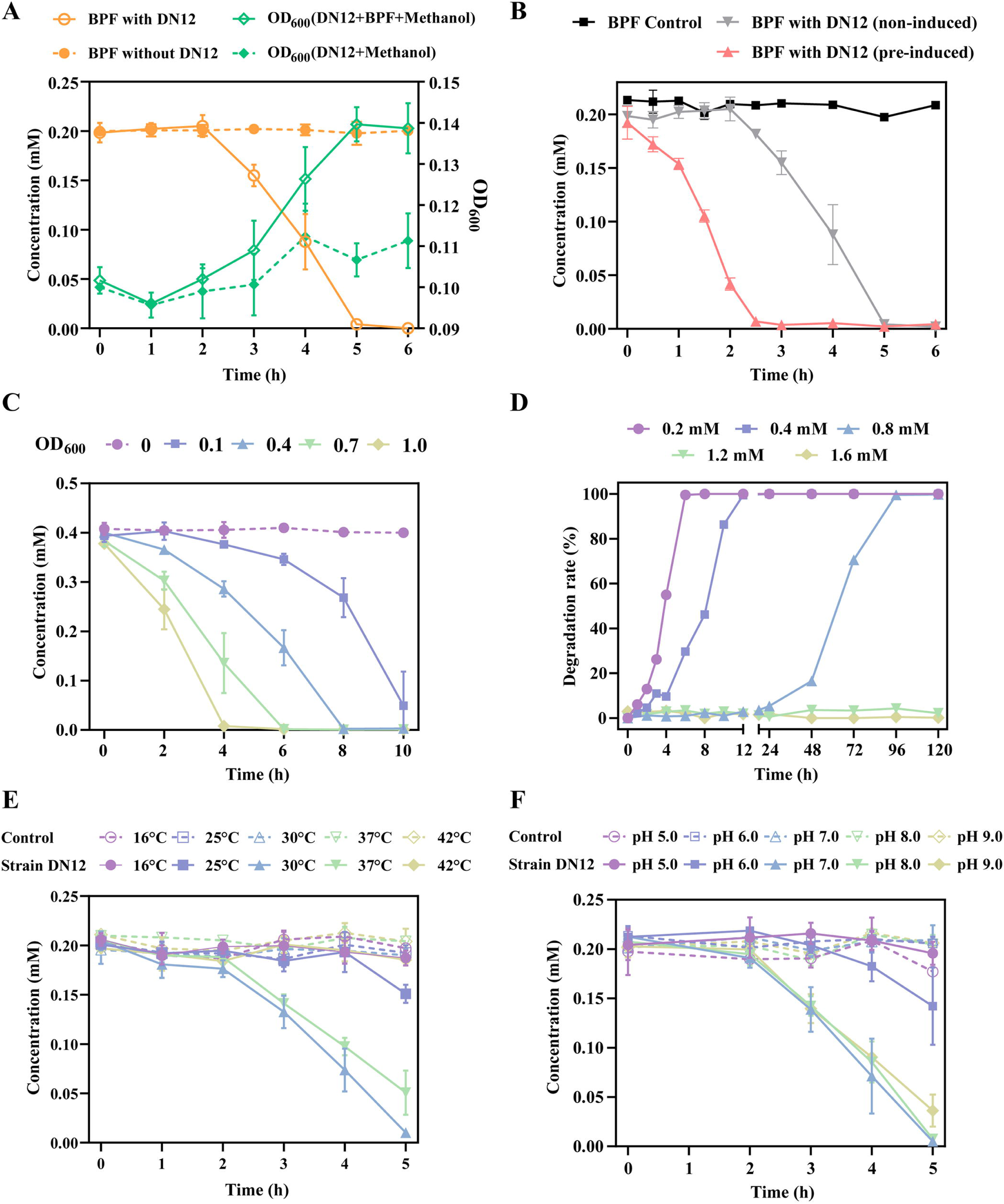
Degradation of BPF by strain DN12. (A) Growth of strain DN12 utilizing BPF as the sole carbon source. BPF concentrations (orange line) and optical density at 600 nm (OD_600_, green line) were monitored over time. (B) Comparative BPF degradation by BPF-induced and uninduced cells of strain DN12. (C) Influence of initial inoculum concentrations on BPF degradation. (D) Impact of initial BPF concentrations on its degradation efficiency. Effects of temperature (E) and pH (F) on BPF degradation. In all panels, the dotted line represents the non-inoculated strain DN12 control under corresponding conditions. Data are presented as mean values ± standard deviations from triplicate experiments.

The degradation rate of BPF by strain DN12 was positively correlated with inoculum density. At an OD_600_ of 1.0, the degradation rate exceeded 99% within 4 h, whereas at an OD_600_ of 0.1, the rate was only 6% within the same period, eventually reaching over 87% after 10 h (Fig. 1C). To further compare BPF degradation efficiency across different inoculum densities, specific degradation activities were calculated. The results showed a significant increase in specific activity with higher initial OD_600_ values, ranging from 3.29 ± 0.99 μmol·OD^-1^·h^-1^ at OD_600_ = 0.1 to 15.52 ± 0.41 μmol·OD^-1^·h^-1^ at OD_600_ = 1.0 (Fig. S3). These data indicate that higher inoculum densities not only accelerated the absolute removal of BPF but also improved the degradation activity per unit of biomass (Fig. 1D). Strain DN12 completely removed 0.2 and 0.4 mM BPF within 12 h, while 0.8 mM BPF was degraded within 48 to 96 h. Higher BPF concentrations (1.2 mM and 1.6 mM) inhibited biodegradation. After 5 d of exposure, CFU-counts of strain DN12 significantly decreased under 1.2 and 1.6 mM BPF compared with the methanol controls (Fig. S4), and HPLC analysis showed no detectable metabolites or substrate depletion at these concentrations (Fig. S5). Both temperature and pH had significant effects on degradation efficiency (Fig. 1E and 1F). Optimal degradation occurred at 30°C, with 0.19 mM BPF degraded within 5 h, corresponding to an average degradation rate of 0.76 µmol·h^-1^. Strain DN12 achieved the highest degradation rates at pH 7.0 and 8.0 (0.78 and 0.77 µmol·h^-1^ within 5 h, respectively), whereas the rates were significantly lower under acidic conditions.

### Identification of the metabolites produced during BPF degradation by strain DN12

High performance liquid chromatography (HPLC) analysis revealed that strain DN12 generated three primary metabolites: M1, M2 and M3 (Fig. S6A). The metabolite M1 was proposed to be DHBP, exhibiting a peak at *m/z* 213.0558 [M-H]^-^ with fragment ions at *m/z* 93.0370 and 143.0520 (Fig. S6B); M2 was identified as HPHB, displaying a peak at *m/z* 229.0507 [M-H]^-^ with a fragment ion at *m/z* 109.0307 (Fig. S6C); M3 was suggested to be 4HB, showing a peak at *m/z* 137.0248 [M-H]^-^ with a fragment ion at *m/z* 93.0368 (Fig. S6D). M1, M2, and M3 had the same retention times, *m/z* values and fragmentation patterns as authentic standards of DHBP, HPHB, and 4HB, respectively.

### Prediction of BPF catabolic genes based on comparative transcriptome analysis

Comparative transcriptome analysis revealed that 73 transcripts, accounting for approximately 1.3% of the total transcripts in strain DN12 (Fig. 2A), were significantly upregulated, with 48 of these exhibiting a greater than 4-fold (log_2_ fold change = 2) increase in transcription in BPF-induced cells compared to uninduced cells (Table S4). In the KEGG pathway analysis, the highest number of enriched genes were associated with *microbial metabolism in diverse environments* (KEGG pathway: map01120) and *benzoate degradation* (KEGG pathway: map00362) (Fig. S7). Within this enriched subset, two duplicated gene clusters, designated as *bpf1* (containing *bpfA1B1*, *bpfC1*, and *bpfD1*) and *bpf2* (containing *bpfA2B2*, *bpfC2*, and *bpfD2*), are hypothesized to be involved in the catabolism of BPF. The encoded enzymes of these two clusters exhibit nearly identical amino acid sequences. Specifically, BpfA1 and BpfA2 share 99% sequence identity, while BpfB1/BpfB2, BpfC1/C2, and BpfD1/D2 are entirely identical (100%). In particular, *orf5019* (*bpfA1*) and *orf5022* (*bpfA2*), located in consecutive but divergently transcribed gene clusters, are predicted to encode FAD-binding oxidoreductases, showing the highest sequence identities to the flavoprotein subunit PchF (43%). PchF, together with the cytochrome *c* subunit PchC, constitutes a two-component *p*-cresol methylhydroxylase (PCMH), which catalyzes the oxidation of *p*-methylphenol (23). Notably, *orf5018* and *orf5021*, located adjacent to *bpfA1* and *bpfA2*, respectively, are annotated as encoding cytochrome *c*. Consequently, *bpfA1B1* and *bpfA2B2* are hypothesized to be responsible for the initial transformation of BPF to DHBP. Both *orf5015* (*bpfC1*) and *orf5026* (*bpfC2*) are annotated as encoding FAD-dependent monooxygenases, with 44% identities to a BVMO from *Pseudomonas putida* ATCC 17453, respectively (24). Therefore, *bpfC1* and *bpfC2* are hypothesized to be involved in the transformation of DHBP to HPHB. Furthermore, *orf5016* (*bpfD1*) and *orf5025* (*bpfD2*) are annotated as encoding α*/*β hydrolases, which are predicted to hydrolyze HPHB to produce 4HB and HQ (Fig. 2B).

**Fig. 2.**
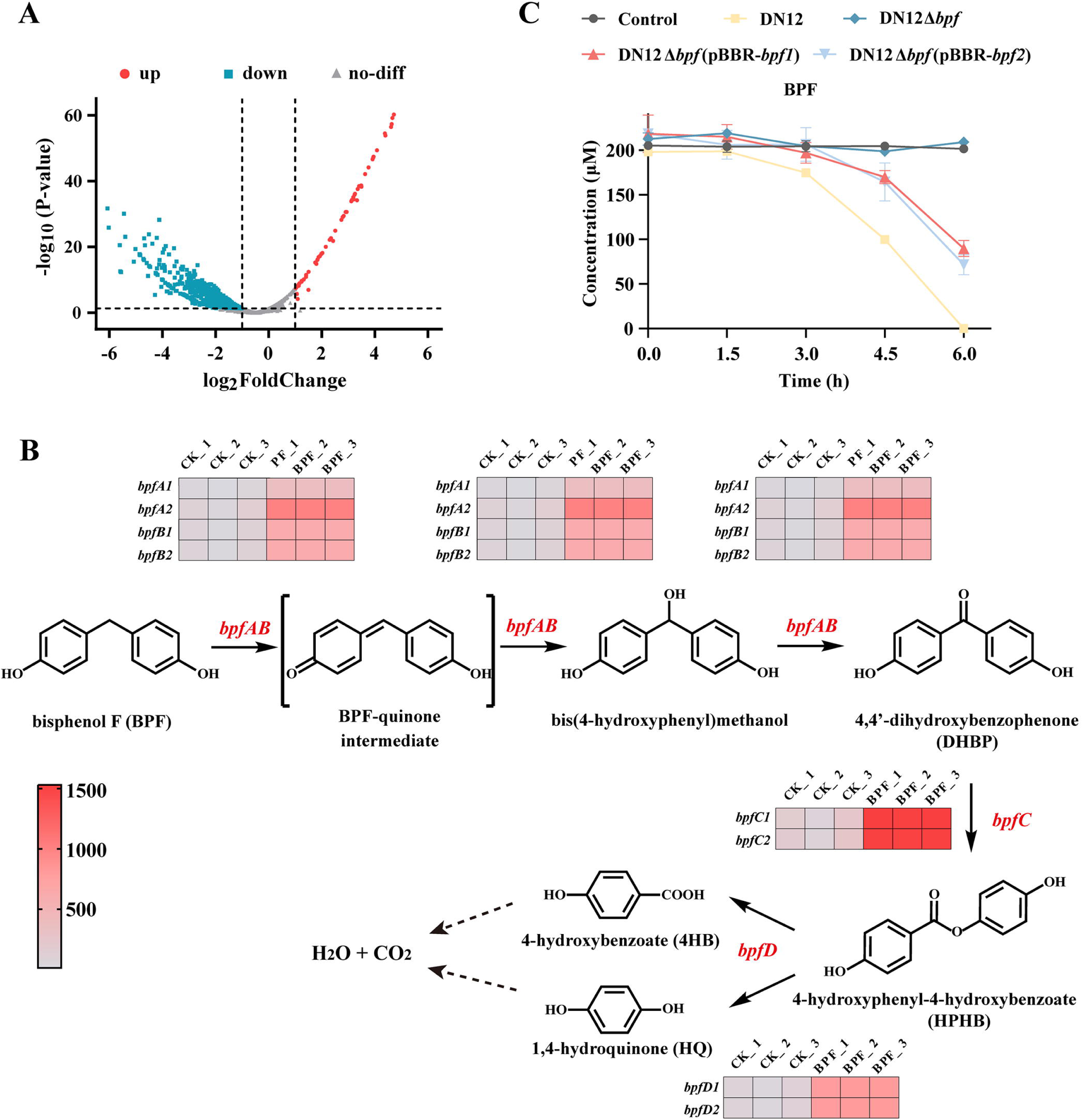
Comparative transcriptome analysis of strain DN12 induced by BPF. (A) Volcano plot of differentially expressed genes following BPF induction. (B) Heatmap of significantly upregulated genes (*bpfABCD*) induced by BPF and their predicted roles in the BPF metabolic pathway, based on functional annotations of the proteins encoded by *bpfABCD*. (C) Degradation of 200 μM BPF in MSM by the wild-type strain DN12, the mutant strain DN12Δ*bpf*, and complemented strains DN12Δ*bpf* (pBBR-*bpf1*) and DN12Δ*bpf* (pBBR-*bpf2*).

To validate the roles of the *bpf* gene cluster in BPF catabolism in strain DN12, a series of mutant and complemented strains were constructed and confirmed through PCR analysis (Fig. S8). Notably, the mutant strain DN12Δ*bpf*, in which *bpf1* (*orf5014-5019*), *orf5020*, *orf5021*, *bpf2* (*orf5022-5027*), *orf5028*, and *orf5029* were deleted, exhibited a complete loss of BPF-degrading capability. In contrast, complemented strains harboring either *bpf1* or *bpf2* regained the ability to degrade BPF (Fig. 2C), confirming that *bpf1* and *bpf2* are essential for BPF catabolism in strain DN12. To elucidate the microbial degradation mechanism of BPF, we investigated the biochemical functions of the proteins encoded by *bpfA1B1*, *bpfC1*, and *bpfD1*.

### The *bpfA1B1* genes encode an oxidoreductase responsible for converting BPF into DHBP

A BLASTP search using the NCBI public database identified 10 proteins with ≥ 38% amino acid sequence homology to BpfA for comparative analysis. BpfA exhibited the highest homology (100%) to the FAD-binding oxidoreductase in *Sphingobium yanoikuyae* SJTF8 (25), though no functional studies have been conducted on this protein. Among characterized proteins, BpfA shared the highest similarity to PCMH (PchF) from *Pseudomonas putida* NCIMB 9869 (43%) (23), followed by PcmI (41%) and PcmJ (39%) from *Geobacter metallireducens* GS-15 (Fig. 3A) (26). PCMH, a well-studied two-component oxidoreductase, has established catalytic mechanisms in both *Pseudomonas* and *Geobacter* species (Fig. S9A).

**Fig. 3.**
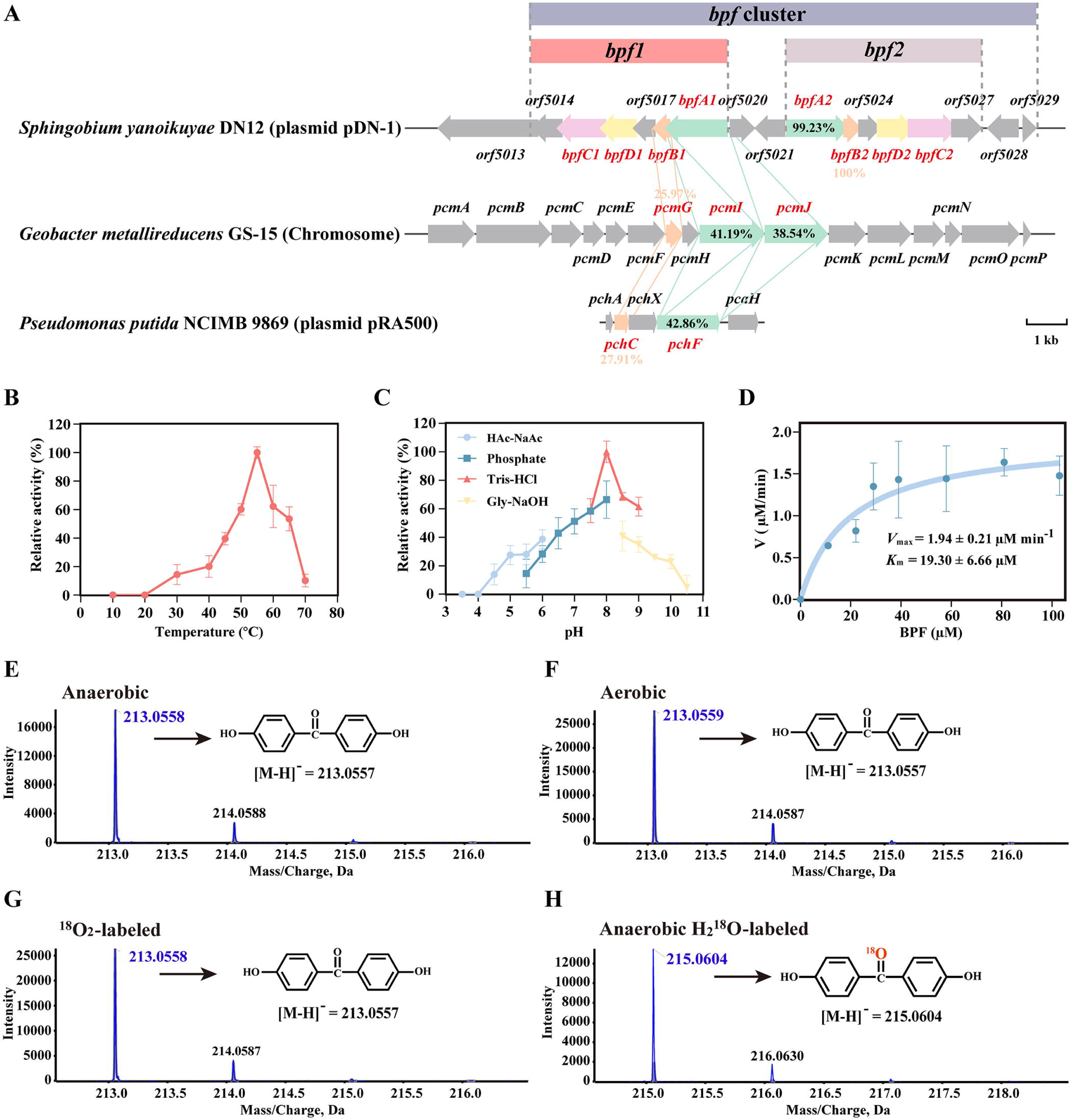
Biochemical characterization of BpfAB. (A) The putative *bpf* degradation gene cluster located on plasmid pDN-1 in strain DN12 and sequence homology of *bpfA* and *bpfB* with the related genes encoding PCMH in *Geobacter metallireducens* GS-15 and *Pseudomonas putida* NCIMB 9869. Effects of temperature (B) and pH (C) on BpfAB enzyme activity. (D) Michaelis-Menten kinetics of BpfAB-mediated BPF oxidation. HRMS analysis of DHBP, the product of BPF transformation by BpfAB under anaerobic (E), aerobic (F), ^18^O_2_-labeled (G), and anaerobic H_2_^18^O-labeled (H) conditions.

BpfAB consists of an FAD-binding oxidoreductase encoded by *bpfA1* (1569 bp) and a *c*-type cytochrome encoded by *bpfB1* (426 bp), with a total of 663 amino acids. Sodium dodecyl sulfate-polyacrylamide gel electrophoresis (SDS-PAGE) analysis showed molecular weights of approximately 58.8 kDa for BpfA and 13.9 kDa for BpfB, both consistent with their theoretical values (Fig. S10). UV-vis analysis of purified BpfAB showed a strong protein absorption peak at 280 nm (Fig. S11A) and FAD-specific peaks at ∼370 and 450 nm (Fig. S11B). Quantitative analysis showed approximately 0.4 µM FAD per 1 µM protein complex. Heme incorporation was minimal: no characteristic Soret peak was observed at 410 nm, but 3,3’,5,5’-tetramethylbenzidine (TMB) staining revealed a faint positive band for BpfAB, absent in BpfA (negative control) yet pronounced in the cytochrome *c* standard (positive control) (Fig. S12). These findings suggest that the majority of the BpfB subunit expressed by *E. coli* exists as apo-cytochromes, with only a minor fraction incorporating heme.

BpfAB displayed optimal activity at pH 8.0 and 55°C, maintaining high activity across pH 6.0-9.0 and 45°C-65°C (Fig. 3B and 3C). Its apparent steady-state kinetic parameters for BPF degradation included a *K*_m_ of 19.30 ± 6.66 μM and *k*_cat_ of 0.79 ± 0.08 s^-1^ (Fig. 3D). To determine the origin of the oxygen atom incorporated into DHBP during the BpfAB-catalyzed reaction, isotope-labeling experiments were performed using H ^18^O and ^18^O_2_, respectively. HRMS analysis showed that the molecular ion peak of DHBP consistently appeared at *m/z* 213.0557 in negative ion mode under anaerobic, aerobic, and ^18^O_2_-labeled conditions (Fig. 3E, 3F, and 3G). In contrast, the DHBP product in the anaerobic H_2_^18^O-labeled group exhibited a +2 *m/z* shift, with the ion peak detected at *m/z* 215.0604 (Fig. 3H), indicating that the hydroxyl oxygen in DHBP was derived from water rather than molecular oxygen. Further metabolite analysis identified bis(4-hydroxyphenyl)methanol as a key intermediate in the transformation of BPF to DHBP (Fig. S13). This intermediate also displayed a +2 *m/z* shift in the H ^18^O-labeled group, suggesting that DHBP formation occurs through multiple sequential steps. In line with the established PCMH model, we propose that this process likely involves the transient formation of a BPF-quinone intermediate, which is subsequently attacked by water molecules to yield hydroxylated products (Fig. 2B). Moreover, the specific activities of BpfAB under anaerobic and aerobic conditions were 0.12 U/mg and 0.14 U/mg, respectively, with no significant difference (*P* > 0.05) (Fig. S14A), further supporting that the hydroxylation step is independent of molecular oxygen. Collectively, these findings demonstrate that in the BpfAB-catalyzed reaction, the oxygen atom incorporated into DHBP originates from water.

To advance our understanding of the phylogeny and habitat distribution of the initiation BPF-degrading enzyme BpfA, we analyzed 333 BpfA-like homologs from prokaryotic genomes retrieved from public datasets. Phylogenetic analysis of 325 genomes revealed that these BpfA-encoding species predominantly belong to the phylum *Pseudomonadota*, accounting for 93% (303 genomes) (Fig. S15). Within this phylum, members of *Burkholderiaceae* (63%) and *Sphingomonadaceae* (12%) were particularly dominant, suggesting that these members have a greater competitive advantage in the face of BPF stress environment. Habitat distribution analysis showed that species encoding BpfA-like proteins are widely distributed across terrestrial and aquatic environments, with 46% associated with soils (Fig. S16). Although BpfA-encoding species in our study originated from aquatic habitats, the significant representation in soils highlights the broader ecological relevance of these organisms. Additionally, genomes of phylogenetically close relatives displayed highly conserved BpfA-like proteins, even when isolated from distinct environments (Fig. 4), reflecting the evolutionary stability of these genes and their vertical inheritance in maintaining functional adaptation across ecological niches (27).

**Fig. 4.**
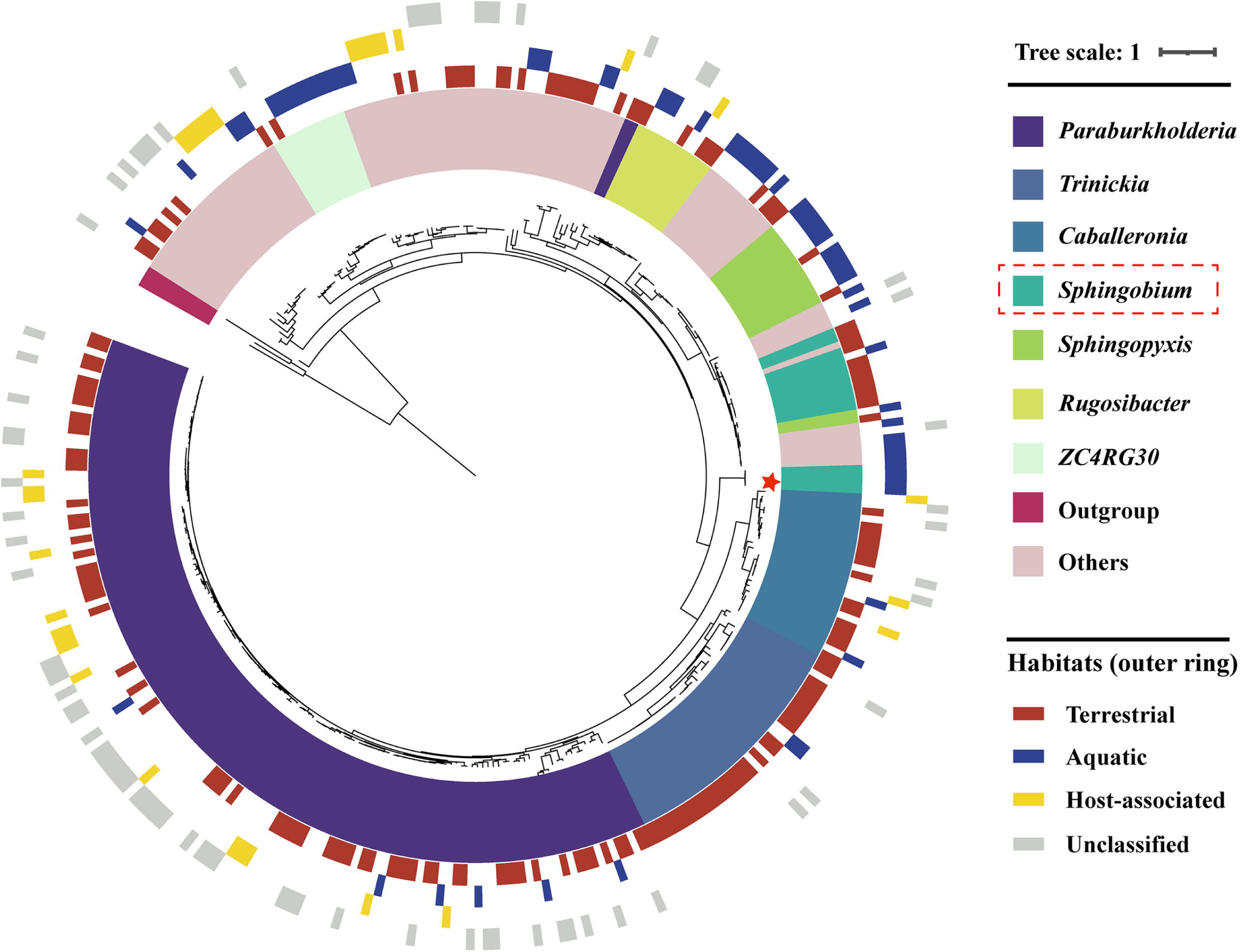
Phylogenetic distribution of BpfA-like homologs based on taxonomy and habitat types. BpfA characterized in this study is marked with a star. Habitats are grouped into four major categories: terrestrial, aquatic, host-associated, and unclassified. Terrestrial habitats mainly include soils and other land-based environments, while aquatic habitats encompass seawater, marine sediments, freshwater, and wastewater.

### The *bpfC1* gene encodes a monooxygenase responsible for converting DHBP into HPHB

BpfC is an FAD-dependent monooxygenase encoded by *bpfC1* (1209 bp), comprising 402 amino acids. SDS-PAGE analysis showed that the size of BpfC was approximately 45.7 kDa (Fig. S10). The optimal activity of BpfC was observed at pH 8.5 and 40°C, with high activity between pH 5.0 and 9.0 and temperatures of 35°C to 50°C (Fig. 5A and 5B). The apparent steady-state kinetic parameters of BpfC for DHBP were *K*_m_ of 21.31 ± 2.88 μM and *k*_cat_ of 2.70 ± 0.13 s^-1^, respectively (Fig. 5C). BpfC catalyzed the conversion of DHBP to HPHB, a typical Baeyer-Villiger rearrangement reaction, where a ketone is oxidized to the corresponding ester. To trace the origin of the oxygen atom incorporated into HPHB, isotope labeling experiments were conducted separately using H ^18^O and ^18^O_2_. HRMS analysis showed that the molecular ion peak of HPHB consistently appeared at *m/z* 229.0506 in negative ion mode under anaerobic, aerobic, and aerobic H ^18^O-labeled conditions (Fig. 5D, 5E, and 5G). In contrast, a distinct +2 *m/z* shift (from 229.0504 to 231.0551) was observed in the ^18^O_2_-labeled group (Fig. 5F), indicating that the oxygen atom incorporated into HPHB originates from molecular oxygen. Time-course assays showed that BpfC catalyzed the progressive formation of HPHB from DHBP under aerobic conditions, whereas only a small amount of HPHB (∼5 μM) was detected within the first 5 min under anaerobic (N_2_-purged) conditions, after which product accumulation ceased (Fig. S17). Additionally, the specific activity of BpfC under aerobic conditions was 0.17 U/mg, significantly higher than that under anaerobic conditions (0.05 U/mg, *P* < 0.05) (Fig. S14B). The weak enzymatic activity observed in the anaerobic group may be attributed to residual oxygen that was not completely removed by nitrogen purging. Collectively, these results demonstrate that the BpfC-catalyzed Baeyer-Villiger oxidation of DHBP is oxygen-dependent.

**Fig. 5.**
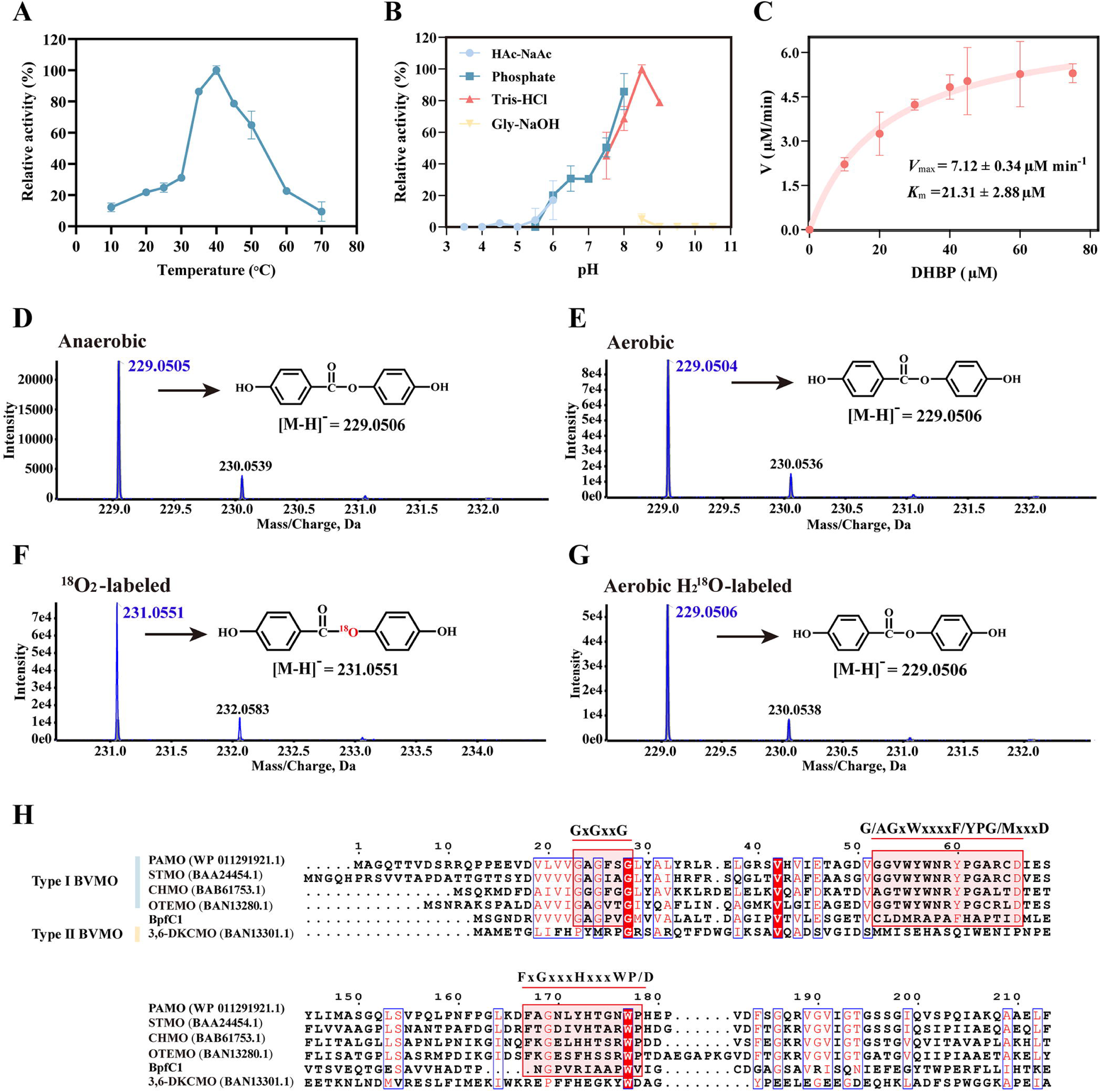
Biochemical characterization of BpfC. Effects of temperature (A) and pH (B) on BpfC enzyme activity. (C) Michaelis-Menten kinetics of BpfC-mediated DHBP oxidation. HRMS analysis of HPHB, the product of DHBP transformation by BpfC under anaerobic (D), aerobic (E), ^18^O_2_-labeled (F), and aerobic H ^18^O-labeled (G) conditions. (H) Amino acid sequence alignment of BpfC with closely related proteins. Conserved domains are marked with red font, and residues proposed to be involved in BpfC substrate specificity are boxed in red.

Until now, three types of BVMOs, Type I (FAD-dependent monooxygenases), Type II (FMN-dependent monooxygenases), and atypical BVMOs (FAD-dependent, but lacks the characteristic sequence of Type I and can catalyze a variety of oxidation reactions) have been characterized (28). Phylogenetic analysis showed that BpfC is a new member of Type I BVMOs (Fig. S9B). Moreover, conserved domain analysis showed that BpfC contained a conserved Rossmann fold FAD-binding domain (GxGxxG/A) and two typical Type I BVMO domains (G/AGxWxxxxF/YPG/MxxxD and FxGxxxHxxxWP/D) (Fig. 5H) (29), further confirming that BpfC is indeed a Type I BVMO.

### The *bpfD1* gene encodes a hydrolase responsible for converting HPHB into 4HB

BpfD is encoded by a putative hydrolase gene *bpfD1* (906 bp), which comprises 301 amino acids. SDS-PAGE analysis showed that BpfD has an approximate molecular weight of 32.9 kDa (Fig. S10). A BLASTP search in the Swiss-Prot database identified seven proteins with ≥ 24% sequence identity to BpfD, all belonging to the α*/*β-hydrolase fold family (Fig. S9C). Enzymatic assays in *vitro* showed that BpfD catalyzes the conversion of HPHB to 4HB and HQ (Fig. 6A and 6B). BpfD exhibited optimal activity at pH 9.0-9.5 and 40°C, and maintained high activity across a broad range of pH values (7.5-10.5) and temperatures (25°C to 50°C) (Fig. 6C and 6D). The apparent steady-state kinetic parameters of BpfD for HPHB were *K*_m_ of 22.54 ± 5.21 μM and *k*_cat_ of 40.24 ± 0.23 s^-1^, respectively (Fig. 6E).

**Fig. 6.**
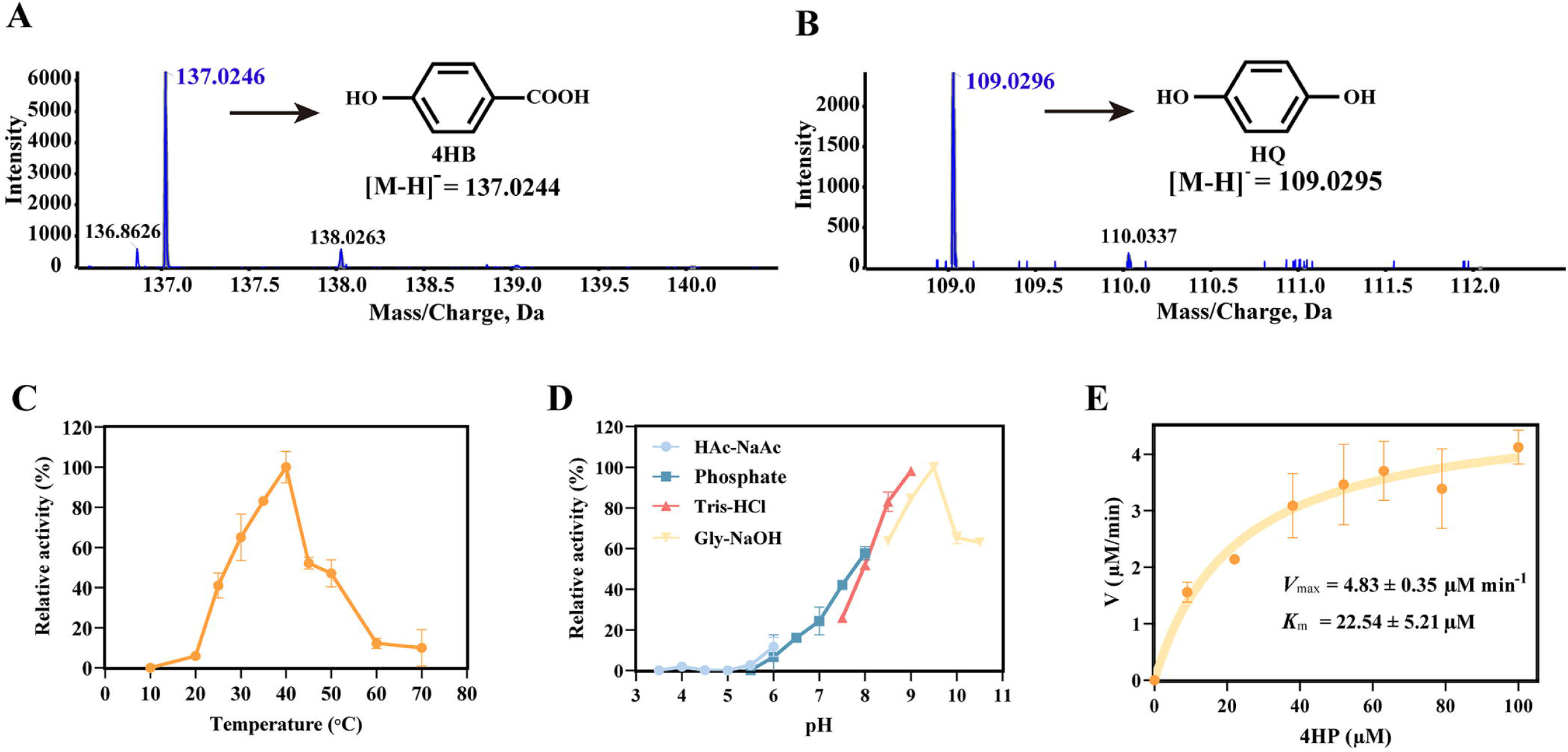
Biochemical characterization of BpfD. HRMS analysis showing that BpfD catalyzes the hydrolysis of HPHB to produce 4HB (A) and HQ (B). Effects of temperature (C) and pH (D) on BpfD enzyme activity. (E) Michaelis-Menten kinetics of BpfD-mediated HPHB hydrolysis.

Sequence and phylogenetic analyses classified BpfD within the α*/*β-hydrolase family. BpfD harbors a conserved GxSxG motif (residues 100-105), characteristic of the catalytic site of this family, as well as the canonical catalytic triad Ser102-Asp247-His275 (Fig. S9D) (30, 31). Collectively, BpfD plays an important role in the BPF metabolic pathway by catalyzing the hydrolysis of a complex compound into simpler central aromatic metabolites, thereby enabling the further mineralization of the parent compound.

## DISCUSSION

To date, the degradation pathway of BPF has been identified in bacteria such as *Pseudomonas* and *Sphingobium*, as well as in white-rot fungi (19–21, 32). Although genetic and biochemical evidence is still lacking, these studies consistently indicate that the symmetrical and stable structure of BPF makes the bridgehead carbon the preferred site for microbial attack, with hydroxylation emerging as an effective degradation pathway. In this study, we identified the same BPF catabolic pathway in strain DN12 and provided molecular-level evidence: BPF is initially converted to DHBP by the two-component oxidoreductase BpfAB; DHBP is then further oxidized to HPHB by the BVMO BpfC; HPHB is hydrolyzed by the hydrolase BpfD to form 4HB and HQ, which ultimately enter the aromatic ring cleavage pathway.

BpfAB is a two-component oxidoreductase composed of an FAD-binding oxidoreductase (BpfA) and a *c*-type cytochrome (BpfB). It catalyzes the hydroxylation of BPF at the bridgehead carbon to DHBP, consistent with the reactions catalyzed by PCMHs (23, 26). Isotope-labeling experiments further confirmed that the oxygen atoms incorporated during BpfAB-catalyzed hydroxylation are derived from water, highlighting a distinct feature compared to canonical oxygenase reactions. In *P. putida* NCIMB 9869/9866, PCMH forms an α*_2_*β*_2_* complex (PchF-PchC) in which electrons are transferred from the substrate to FAD and subsequently to the heme cofactor, representing the typical electron-transfer mechanism of PCMH-like enzyme (23, 33–35).

Our sequence analysis supports a non-covalent FAD-binding mode in BpfA, distinguishing it from classical PCMHs. Alignment revealed the absence of the conserved tyrosine residue (e.g., Tyr384 in PchF of *P. putida*, Tyr394 in PcmI of *G. metallireducens*) that mediates covalent FAD attachment. Instead, BpfA harbors a glycine-rich motif (GLDGYR, residues 38-43) characteristic of the phosphate-binding loop within a Rossmann fold, a canonical domain for non-covalent dinucleotide cofactor binding (36) (Fig. S18). The presence of this Rossmann signature, together with the absence of the covalent linkage site, indicates that BpfA adopts a “loose binding” strategy for FAD. In parallel, the limited heme incorporation into BpfB is consistent with the limited efficiency of cytochrome *c* maturation in *E. coli*, a well-recognized bottleneck for heterologous expression of *c*-type cytochromes that depends on the host’s cytochrome *c* maturation system, which often fails to efficiently process foreign proteins (37).

Sequence and localization analyses further suggest that BpfAB shares structural similarity with these homologous systems. SignalP 6.0 predicted a Sec/SPII signal peptide in BpfB but not in BpfA, a pattern also observed in PchCF (*P. putida*) and PcmIG (*G. metallireducens*). Previous studies confirmed that PchCF is a periplasmic complex (35), supporting the hypothesis that BpfAB is also localized to the periplasm, guided by the signal peptide of BpfB. In contrast to the *Geobacter* Pcm system, no homologs of *pcmCDEF* (encoding a membrane-anchoring complex) were detected around the *bpf* cluster. This resembles the PchCF system, which also lacks PcmCDEF and instead relies on soluble periplasmic electron carriers such as azurin (38). Collectively, these findings suggest that BpfAB likely functions as a soluble periplasmic complex in its native host. Importantly, the identification of BpfAB demonstrates that PCMH-like enzymes are not restricted to small aromatic substrates such as *p*-cresol but can also hydroxylate structurally complex synthetic bisphenols, expanding the catalytic repertoire of this enzyme family and providing a molecular basis for microbial strategies to mitigate bisphenol pollution.

BpfC is a typical BVMO, which catalyzes the oxidation reaction of DHBP to HPHB. BVMOs have been widely used in biocatalysis field, and numerous studies have demonstrated their ability to catalyze the transformation of various substrates, including aromatic and linear ketones, aldehydes, bicyclic ketones, and steroids (39, 40). In established Baeyer-Villiger reactions, oxygen atom is typically inserted at the substituent with the highest electron-donating potential on the ketone group, showcasing precise substrate and regioselectivity (41). Notably, most BVMOs catalyze reactions with cyclic ketones or asymmetric structures (42–45). In this study, we discovered that BpfC catalyzes the oxidation of a symmetric substrate, DHBP, which features identical structures on both sides of the ketone group, thereby expanding the substrate scope of BVMOs.

In this study, we conducted detailed physiological and biochemical analyses of BpfAB, BpfC, and BpfD. Specifically, it comprehensively reveals the biochemical roles of three degradation enzymes in the key three-step oxidation-oxidation-hydrolysis process during BPF conversion. This work fills a gap in the molecular mechanisms of BPF degradation by microorganisms.

## MATERIALS AND METHODS

### Chemical reagents and media

BPF, BPA, BPS, TBBPA, and TBBPS were purchased from Aladdin Biotechnology (Shanghai, China). DHBP, HPHB, 4HB, BPB, BPE, and TDP were purchased from Bide Pharmatech (Shanghai, China). H_2_ ^18^O (98% atom ^18^O) was purchased from AngelChem (Shanghai, China), and chromatographic-grade methanol was purchased from Merck (Germany). All other reagents used in this research were analytical grade. LB liquid medium, MSM, and R2A medium were prepared as previously reported (46).

### BPF-degrading bacteria isolation from a river sediment

Sediments collected from a river in Anhui Province, China, were inoculated into 100 mL of MSM (v/v, 1/10) supplemented with 0.1 mM BPF and incubated at 180 rpm and 30°C for 7 days. The initial culture was subsequently transferred (v/v, 1/10) into MSM containing 0.2 mM BPF for multiple enrichment cycles. After three rounds of cultivation, the enriched culture was serially diluted and plated onto 1/5 LB and 1/2 R2A agar supplemented with 0.2 mM BPF for cultivation. Single colony was selected for culture, BPF degradation efficiency of each bacterial isolate was analyzed using HPLC. Through repeated cycles of isolation and testing, a bacterial strain designated DN12, with efficient BPF-degrading capability, was obtained from the enrichment cultures. The morphology of strain DN12 was analyzed using scanning electron microscopy (SEM, SU8010, Hitachi), and its phylogenetic position was determined through a neighbor-joining phylogenetic tree constructed based on 16S rRNA gene sequences using MEGA 7.0 software.

### Biodegradation assay by strain DN12

Cells of strain DN12 were inoculated into 100 mL of LB broth and cultured at 30°C and 180 rpm until reaching the exponential growth phase. The cells were then harvested by centrifugation at 6000 rpm for 8 min, washed twice with MSM, and resuspended in MSM to prepare the seed cells. These seed cells were inoculated into 20 mL of MSM (initial OD_600_ = 0.1) supplemented with 0.2 mM BPF and incubated at 30°C and 180 rpm. Subsequently, 0.5 mL of culture was sampled at regular intervals (1 h), centrifuged at 12000 rpm for 5 min, and detected by HPLC. Using a similar approach, the BPF-degrading capacity of strain DN12 was evaluated under varying conditions, including temperatures (16, 25, 30, 37, and 42°C), pH levels (5.0, 6.0, 7.0, 8.0, and 9.0), initial inoculum densities (OD_600_ of 0.1, 0.4, 0.7, and 1.0), and initial BPF concentrations (0.2, 0.4, 0.8, 1.2 and 1.6 mM). All experiments were performed in triplicate.

### Whole genome sequencing and comparative transcriptomic analysis

Strain DN12 cells were collected after incubation at 30 for 12 h, and genomic DNA was extracted using the Invitrogen PureLink® Genomic DNA kit. Whole-genome sequencing of strain DN12 was performed by Shanghai Biozeron Biotechnology Co., Ltd. (Shanghai, China), using the Illumina NovaSeq 6000 paltform (PE150 Mode) and Pacific Biosciences Sequel IIe technology (PacBio).

To identify the genes and enzymes involved in BPF metabolism, a comparative transcriptomic analysis was performed based on phenotypic differences observed in strain DN12 under substrate induction. Strain DN12 cells were initially cultured in LB medium until the exponential growth stage; the culture was then divided into two groups: the induced group was supplemented with 0.2 mM BPF, while the control group was supplemented with an equivalent volume of methanol. After co-culturing for 3 h, the BPF-induced and uninduced cells were harvested for induced degradation experiment and RNA-seq. Differential gene expression analysis was conducted using EdgeR (https://bioconductor.org/packages/release/bioc/html/edgeR.html), with functional enrichment analyses performed using Gene Ontology (GO) and Kyoto Encyclopedia of Genes and Genomes (KEGG) annotations.

### Cloning and functional verification of the *bpf* gene cluster

To confirm the function of the *bpf* gene cluster, gene knockout and complementation experiments were conducted. Homologous arms of 800 bp upstream and downstream of the gene cluster *bpf* were amplified from genomic DNA from strain DN12 using primers Δ*bpf*_upF/R and Δ*bpf*_downF/R (Table 1). These fragments were assembled by overlap extension PCR and inserted into the suicide vector pJQ200SK to construct pJQ-*bpf*, which was then introduced into strain DN12 via triparental mating with the helper strain *E. coli* HB101 (pRK2013). Mutant strains were isolated through streptomycin-positive and sucrose-negative selection and designated DN12Δ*bpf* (Table 2).

**Table 1.**
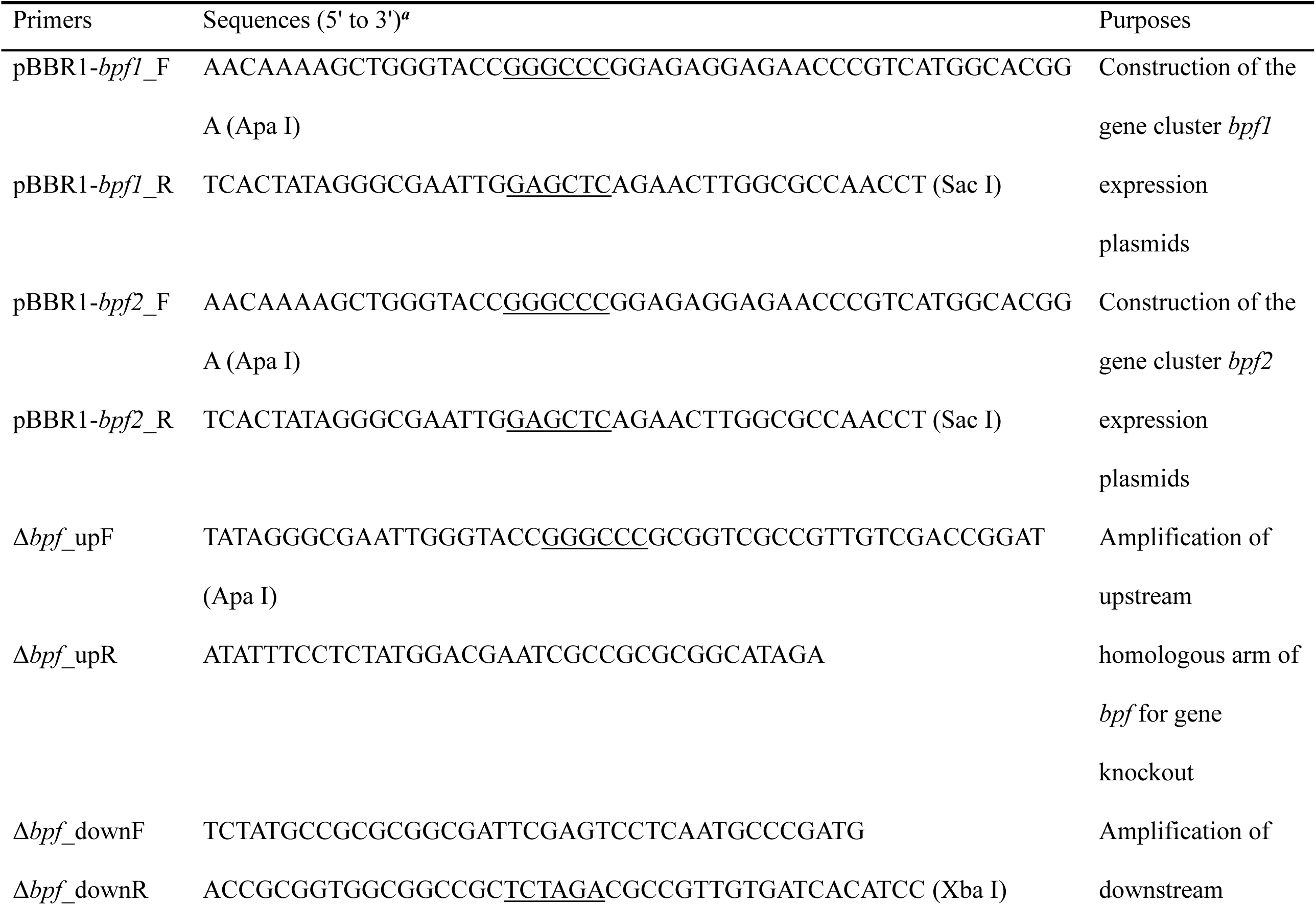
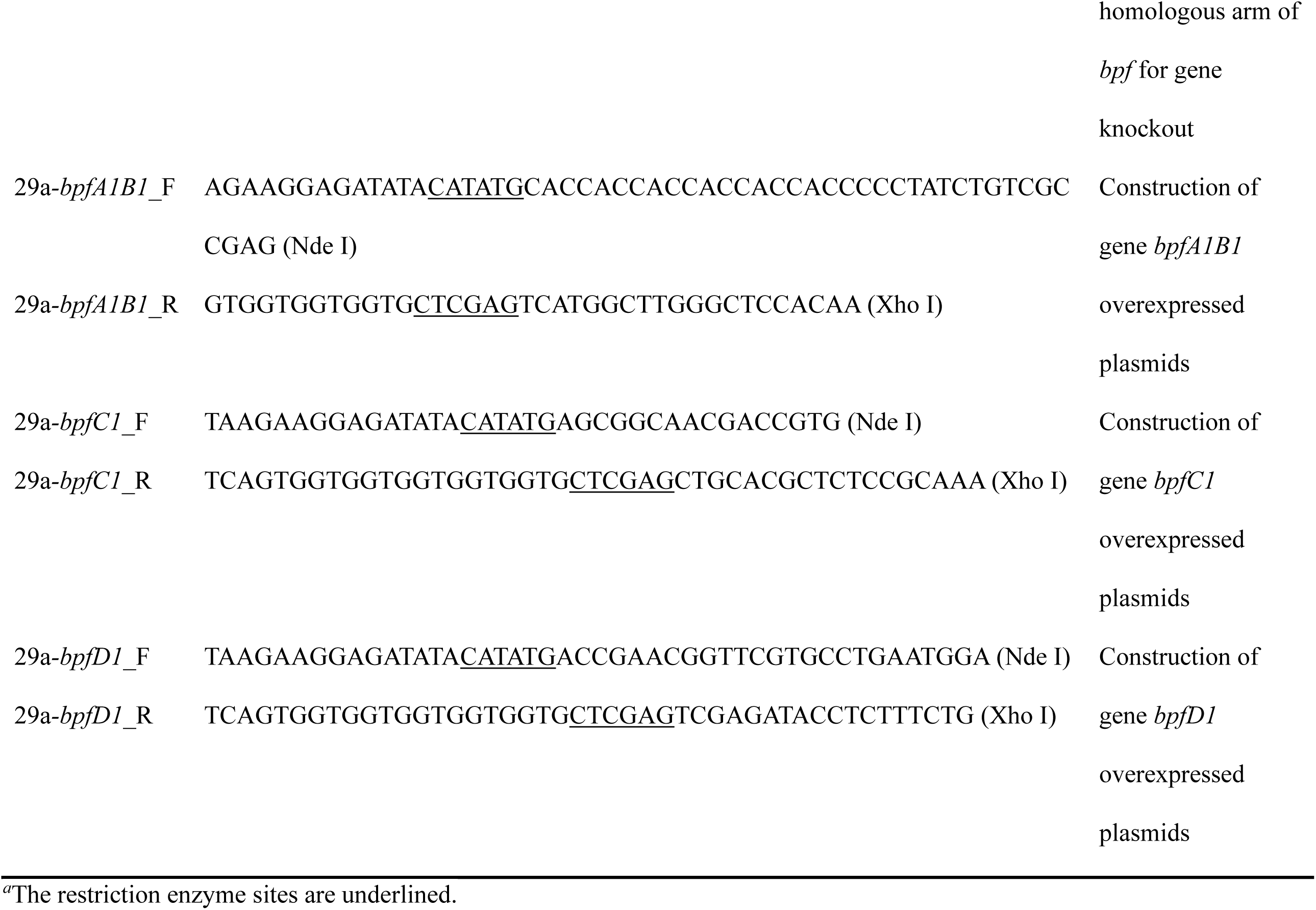
Primers used in this study.

**Table 2.**
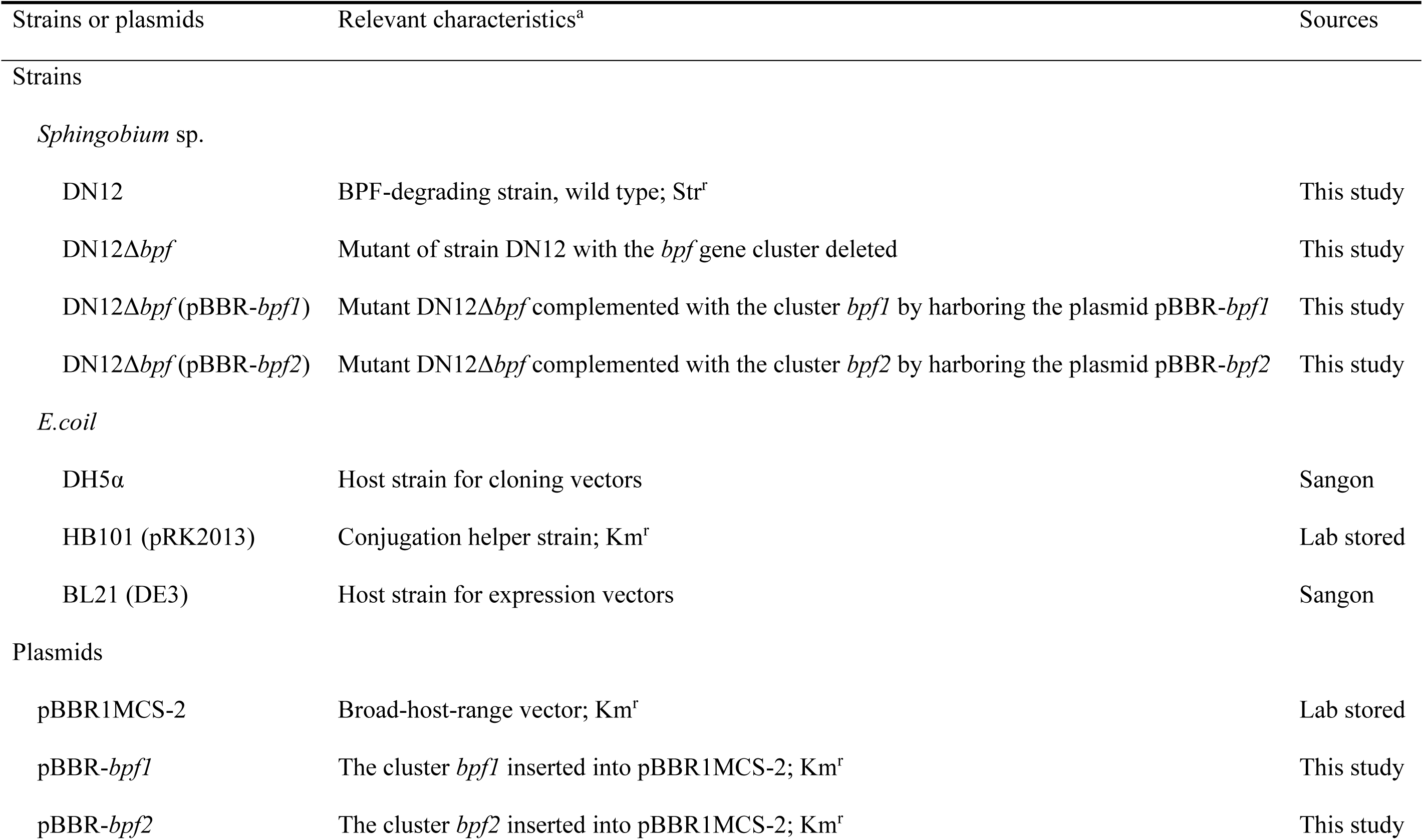
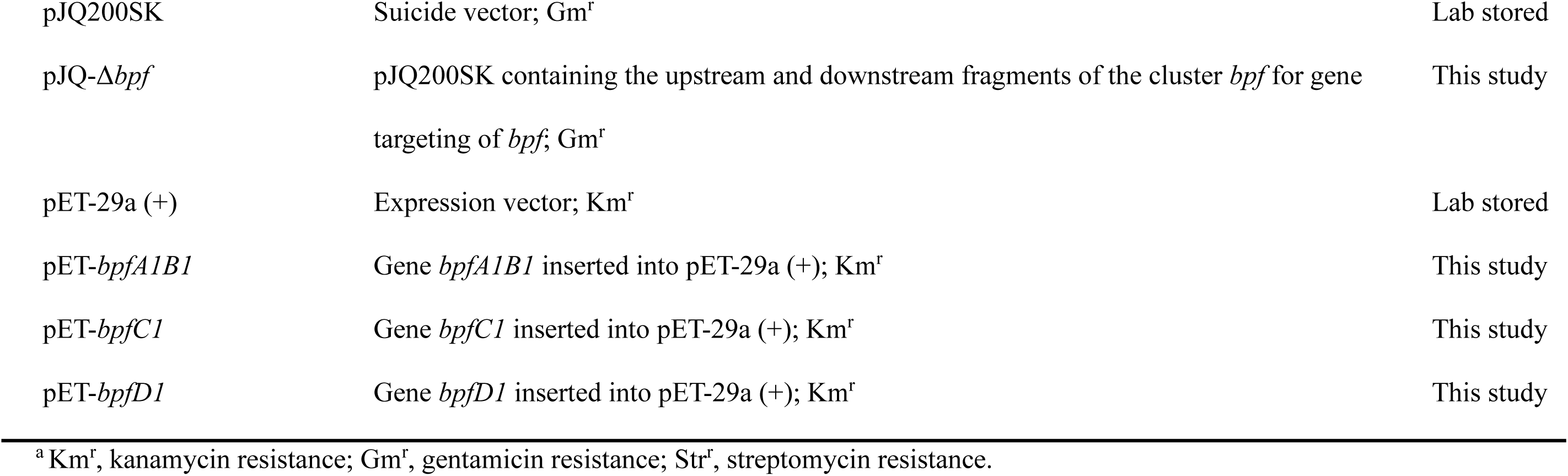
Bacterial strains and plasmids used in this study.

The clusters *bpf1* and *bpf2* were amplified using the primers pBBR1-*bpf1*_F/R, and pBBR1-*bpf2*_F/R (Table 1), respectively, and cloned into the broad-host-range plasmid pBBR1MCS-2 to generate pBBR-*bpf1* and pBBR-*bpf2*. Subsequently, these recombinant plasmids were introduced into the mutant strain DN12Δ*bpf* via triparental mating to obtain complementary strains DN12Δ*bpf* (pBBR-*bpf1*) and DN12Δ*bpf* (pBBR-*bpf2)*.

### Protein expression and purification

The BPF degradation genes *bpfA1B1*, *bpfC1* and *bpfD1* were amplified from the gene cluster *bpf1* using primers 29a-*bpfA1B1*_F/R, 29a-*bpfC1*_F/R, and 29a-*bpfD1*_F/R (Table 1), respectively, and then cloned into the overexpression vector pET29a (+) to generate pET-*bpfA1B1*, pET-*bpfC1*, and pET-*bpfD1* (Table 2). Recombinant plasmids were transformed into *E. coli* BL21(DE3) and cultured in LB medium supplemented with 50 mg·L^-1^ kanamycin. Protein expression was induced with 0.2 mM IPTG, after which cells were harvested and disrupted by ultrasonication to obtain crude extract. His-tagged proteins were purified by nickel affinity chromatography (Ni-NTA) at 4, followed by gradient elution with imidazole (47). The purified target protein was analyzed by SDS-PAGE and subsequently dialyzed against two liters of PBS (8.0 g NaCl, 0.2 g KCl, 1.42 g Na_2_HPO_4_, 0.24 g KH_2_PO_4_ per liter of water, pH 7.2). Protein concentration was determined by the Bradford method using bovine serum albumin as the standard (48).

### Determination of FAD and heme content in BpfAB

UV-vis absorption spectra of purified BpfAB were recorded using a spectrophotometer (Evolution One Plus, Thermo Scientific) in the range of 250-600 nm. FAD content was quantified by comparing the absorbance at 450 nm with a standard curve generated from authentic FAD standards prepared in the same buffer. For heme detection, purified proteins were separated on 12% SDS–PAGE gels and subsequently analyzed using the TMB heme-staining method as described previously (49). Briefly, gels were incubated in freshly prepared TMB staining solution (0.25 mg·mL^-1^ TMB and 0.03% H_2_O_2_ in 0.25 M sodium acetate buffer, pH 5.0) until bands appeared, then rinsed with distilled water to stop the reaction. Horse heart cytochrome *c* (Sigma-Aldrich) and purified BpfA were used as positive and negative controls, respectively.

### Enzymatic activity assay

Three standard enzymatic reaction systems were established to characterize enzyme activity: (1) BpfAB activity was measured in 500 µL of PBS containing 0.1 mM flavin adenine dinucleotide (FAD), 0.2 mM phenazine methosulfate (PMS), 14–140 nM BpfAB, and 0.2 mM BPF; (2) BpfC activity was measured in 500 µL of PBS containing 0.02 mM FAD, 0.4 mM nicotinamide adenine dinucleotide phosphate (NADPH), 21–218 nM BpfC, and 0.2 mM DHBP; (3) BpfD activity was measured in 500 µL of PBS containing 0.021 µM BpfD and 0.2 mM HPHB. Reactions were terminated by boiling for 5 min, followed by centrifugation at 16,000 × g for 10 min. The supernatants were filtered through a 0.22 µm membrane and analyzed by HPLC to quantify the residual substrates. The optimal temperature and pH for enzyme activity were determined using the standard enzymatic reaction system at varying temperatures (10-70°C) and pH values (3.5-10.5) adjusted with different buffers. The kinetic parameters (*K*_m_ and *k*_cat_) of BpfAB, BpfC, and BpfD were determined under their respective optimal reaction temperature and pH conditions. Nonlinear regression analysis of the Michaelis-Menten equation was performed using GraphPad Prism software, and the fitting errors were calculated accordingly. One unit of enzymatic activity was defined as the amount of enzyme required to convert 1 µmol of substrate per minute. All enzymatic assays were independently repeated three times.

To investigate the oxygen dependency of BpfAB and BpfC, enzymatic reactions were conducted under both anaerobic and aerobic conditions, coupled with isotope labeling using ^18^O_2_ and H ^18^O. Anaerobic reactions were performed in an anaerobic chamber (COY-7000220A, COY Laboratory Products Inc.). Prior to use, all components required for the enzymatic reaction—including the purified enzyme solution, cofactors, and PBS—were placed in serum bottles and deoxygenated by purging with high-purity nitrogen gas (99.99%) for 30 min in an ice-water bath. The specific activities of BpfAB and BpfC were then measured under aerobic and anaerobic conditions.

### Phylogenetic analysis

The amino acid sequences of BpfA, BpfC, and BpfD homologs were constructed using the BLASTP available at the NCBI database (https://blast.ncbi.nlm.nih.gov/Blast.cgi). For evolutionary analysis, the aligned protein sequences were subjected to phylogenetic tree construction using the MEGA 7.0 software. The neighbor-joining algorithm was employed, with bootstrap replicates set at 1000 to assess the statistical reliability of the tree topology. Conserved domains and sequence motifs within the aligned proteins were analyzed using ClustalW (https://www.genome.jp/tools-bin/clustalw) to generate multiple sequence alignments. Structural and motif visualization was performed using ESPript 3.0 (https://espript.ibcp.fr/ESPript/cgi-bin/ESPript.cgi), enabling clear annotation of conserved residues and secondary structure elements (50).

### Identification of BpfA-like in prokaryotic databases

The Prokaryotic Genome dataset was collected from the published literature the (n = 200532) and Integrated Microbial Genomes & Microbiomes (IMG) (n = 157560). The ORFs of genomes from the datasets mentioned above were predicted using Prodigal (v2.6.3) with the parameter ‘-p single’. The resulting protein sequences were aligned to BpfA using DIAMOND (identity≥50%, query coverage>75%, E-value<1×10^−10^) for explore more diverse BpfA homologs. Genome taxonomy was determined by with GTDB-Tk (v2.3.3) using ‘classify_wf’ function according to release 214 database, and phylogenetic relationship were inferred by with GTDB-Tk using the ‘identify’ and ‘align’ function. A maximum likelihood tree was constructed with FastTree (version 2.1.10). BpfA homologs sequences were aligned with MAFFT L-INS-i v7.407, trimmed using TRIMAL 1.2rev59 (settings:-automated1). Protein phylogenetic tree was generated with IQ-TREE v2.1.3 (settings:-m Q.pfam+R7-B 1000). All trees were visualized by iTOL online software.

### Analytical methods

The cultures and enzyme assay samples were first mixed with an equal volume of methanol and then centrifuged (12,000 rpm) for 5 min. The supernatant was filtered through a membrane (0.22 µm pore size) and qualitatively and quantitatively determined using U3000 HPLC system (Thermo Fisher Scientific Inc., Germany) equipped with an Agilent ZORBAX StableBond C18 column (4.6 × 250 mm, 5 µm). The mobile phase consisted of methanol and 0.5% aqueous acetic acid (60:40, v/v) at a flow rate of 0.75 mL·min^-1^. The column was maintained at 35°C during analysis. To further identify the metabolites BPF, a Triple TOF 5600 HRMS (AB Sciex, Framingham, MA, USA) equipped with a Turbo V probe was used. Electrospray ionization was conducted in negative polarity mode.

### Data availability

The complete genome sequence of *Sphingobium yanoikuyae* DN12 has been deposited in the GenBank database under accession numbers CP173717, CP173718, CP173719, and CP173720. Additionally, the transcriptome data for *Sphingobium yanoikuyae* DN12 is available in the GenBank database under bioproject number PRJNA1181554.

## SUPPLEMENTAL MATERIAL

Figures S1-S18; Tables S1-S4.

## ACKNOWLEDGEMENTS

We gratefully acknowledge funding from the grant of the National Key R&D Program of China (2024YFD1502400), the National Natural Science Foundation of China (32370126 and 32070091), and the Postgraduate Research & Practice Innovation Program of Jiangsu Province (KYCX25_1044 and KYCX24_0922).

## REFERENCES

1. Su C, Cui Y, Liu D, Zhang H, Baninla Y. 2020. Endocrine disrupting compounds, pharmaceuticals and personal care products in the aquatic environment of China: Which chemicals are the prioritized ones? Sci Total Environ 720:137652.

2. Xing J, Zhang S, Zhang M, Hou J. 2022. A critical review of presence, removal and potential impacts of endocrine disruptors bisphenol A. Comp Biochem Physiol C Toxicol Pharmacol 254:109275.

3. Algonaiman R, Almutairi AS, Al Zhrani MM, Barakat H. 2023. Effects of Prenatal Exposure to Bisphenol A Substitutes, Bisphenol S and Bisphenol F, on Offspring’s Health: Evidence from Epidemiological and Experimental Studies. Biomolecules 2023, 13(11): 1616.

4. Cimmino I, Fiory F, Perruolo G, Miele C, Beguinot F, Formisano P, Oriente F. 2020. Potential Mechanisms of Bisphenol A (BPA) Contributing to Human Disease. Int J Mol Sci 21.

5. Siracusa JS, Yin L, Measel E, Liang S, Yu X. 2018. Effects of bisphenol A and its analogs on reproductive health: A mini review. Reprod Toxicol 79:96–123.

6. Eladak S, Grisin T, Moison D, Guerquin MJ, N’Tumba-Byn T, Pozzi-Gaudin S, Benachi A, Livera G, Rouiller-Fabre V, Habert R. 2015. A new chapter in the bisphenol A story: bisphenol S and bisphenol F are not safe alternatives to this compound. Fertil Steril 103:11–21.

7. Ye X, Wong LY, Kramer J, Zhou X, Jia T, Calafat AM. 2015. Urinary Concentrations of Bisphenol A and Three Other Bisphenols in Convenience Samples of U.S. Adults during 2000-2014. Environ Sci Technol 49:11834–9.

8. Liu J, Zhang L, Lu G, Jiang R, Yan Z, Li Y. 2021. Occurrence, toxicity and ecological risk of Bisphenol A analogues in aquatic environment - A review. Ecotoxicol Environ Saf 208:111481.

9. Bornehag CG, Engdahl E, Unenge Hallerbäck M, Wikström S, Lindh C, Rüegg J, Tanner E, Gennings C. 2021. Prenatal exposure to bisphenols and cognitive function in children at 7 years of age in the Swedish SELMA study. Environ Int 150:106433.

10. Gu J, Li L, Yin X, Liang M, Zhu Y, Guo M, Zhou L, Fan D, Shi L, Ji G. 2022. Long-term exposure of zebrafish to bisphenol F: Adverse effects on parental reproduction and offspring neurodevelopment. Aquat Toxicol 248:106190.

11. Moreman J, Lee O, Trznadel M, David A, Kudoh T, Tyler CR. 2017. Acute Toxicity, Teratogenic, and Estrogenic Effects of Bisphenol A and Its Alternative Replacements Bisphenol S, Bisphenol F, and Bisphenol AF in Zebrafish Embryo-Larvae. Environ Sci Technol 51:12796–12805.

12. Mu X, Liu J, Wang H, Yuan L, Wang C, Li Y, Qiu J. 2022. Bisphenol F Impaired Zebrafish Cognitive Ability through Inducing Neural Cell Heterogeneous Responses. Environ Sci Technol 56:8528–8540.

13. Yuan L, Qian L, Qian Y, Liu J, Yang K, Huang Y, Wang C, Li Y, Mu X. 2019. Bisphenol F-Induced Neurotoxicity toward *Zebrafish* Embryos. Environ Sci Technol 53:14638–14648.

14. Han Y, Hu LX, Liu T, Liu J, Wang YQ, Zhao JH, Liu YS, Zhao JL, Ying GG. 2022. Non-target, suspect and target screening of chemicals of emerging concern in landfill leachates and groundwater in Guangzhou, South China. Sci Total Environ 837:155705.

15. Rodríguez-Hernández JA, Araújo RG, López-Pacheco IY, Rodas-Zuluaga LI, González-González RB, Parra-Arroyo L, Sosa-Hernández JE, Melchor-Martínez EM, Martínez-Ruiz M, Barceló D. 2022. Environmental persistence, detection, and mitigation of endocrine disrupting contaminants in wastewater treatment plants–a review with a focus on tertiary treatment technologies. Environmental Science: Advances 1:680–704.

16. Tang Z, Liu ZH, Wang H, Wan YP, Dang Z, Guo PR, Song YM, Chen S. 2023. Twelve natural estrogens and ten bisphenol analogues in eight drinking water treatment plants: Analytical method, their occurrence and risk evaluation. Water Res 243:120310.

17. Mirza A, Kumar A, Singh G, Arya SK, Bhalla A, Singh J. 2024. Microbial Technology: Tools for Waste Management; Environmental Sustainability and Environmental Safety, p 41-70, Microbial Applications for Environmental Sustainability. Springer.

18. Lu H, Weng Z, Wei H, Zhou J, Wang J, Liu G, Guo W. 2017. Simultaneous bisphenol F degradation, heterotrophic nitrification and aerobic denitrification by a bacterial consortium. Journal of Chemical Technology & Biotechnology 92:854–860.

19. Inoue D, Hara S, Kashihara M, Murai Y, Danzl E, Sei K, Tsunoi S, Fujita M, Ike M. 2008. Degradation of Bis(4-Hydroxyphenyl)methane (bisphenol F) by *Sphingobium yanoikuyae* strain FM-2 isolated from river water. Appl Environ Microbiol 74:352–8.

20. Toyama T, Ojima T, Tanaka Y, Mori K, Morikawa M. 2013. Sustainable biodegradation of phenolic endocrine-disrupting chemicals by *Phragmites australis*-rhizosphere bacteria association. Water Sci Technol 68:522–9.

21. Toyama T, Sato Y, Inoue D, Sei K, Chang YC, Kikuchi S, Ike M. 2009. Biodegradation of bisphenol A and bisphenol F in the rhizosphere sediment of *Phragmites australis*. J Biosci Bioeng 108:147–50.

22. Zühlke M-K, Schlüter R, Henning A-K, Lipka M, Mikolasch A, Schumann P, Giersberg M, Kunze G, Schauer F. 2016. A novel mechanism of conjugate formation of bisphenol A and its analogues by *Bacillus amyloliquefaciens*: detoxification and reduction of estrogenicity of bisphenols. International Biodeterioration & Biodegradation 109:165–173.

23. Kim J, Fuller JH, Cecchini G, McIntire WS. 1994. Cloning, sequencing, and expression of the structural genes for the cytochrome and flavoprotein subunits of *p*-cresol methylhydroxylase from two strains of *Pseudomonas putida*. Journal of Bacteriology 176:6349–6361.

24. Leisch H, Shi R, Grosse S, Morley K, Bergeron H, Cygler M, Iwaki H, Hasegawa Y, Lau PC. 2012. Cloning, baeyer-villiger biooxidations, and structures of the camphor pathway 2-oxo-δ3-4, 5, 5-trimethylcyclopentenylacetyl-coenzyme a monooxygenase of *Pseudomonas putida* ATCC 17453. Applied and Environmental Microbiology 78:2200–2212.

25. Yin C, Xiong W, Qiu H, Peng W, Deng Z, Lin S, Liang R. 2020. Characterization of the phenanthrene-degrading *Sphingobium yanoikuyae* SJTF8 in heavy metal co-existing liquid medium and analysis of its metabolic pathway. Microorganisms 8:946.

26. Johannes Jr, Bluschke A, Jehmlich N, von Bergen M, Boll M. 2008. Purification and characterization of active-site components of the putative *p*-cresol methylhydroxylase membrane complex from *Geobacter metallireducens*. Journal of Bacteriology 190:6493–6500.

27. Wiedenbeck J, Cohan FM. 2011. Origins of bacterial diversity through horizontal genetic transfer and adaptation to new ecological niches. FEMS Microbiology Reviews 35:957–976.

28. Tolmie C, Smit MS, Opperman DJ. 2019. Native roles of Baeyer–Villiger monooxygenases in the microbial metabolism of natural compounds. Natural Product Reports 36:326–353.

29. Ceccoli RD, Bianchi DA, Fink MJ, Mihovilovic MD, Rial DV. 2017. Cloning and characterization of the Type I Baeyer–Villiger monooxygenase from *Leptospira biflexa*. AMB Express 7:1–13.

30. De Schrijver A, Nagy I, Schoofs G, Proost P, Vanderleyden J, Van Pée K, De Mot R. 1997. Thiocarbamate herbicide-inducible nonheme haloperoxidase of *Rhodococcus erythropolis* NI86/21. Applied and Environmental Microbiology 63:1911–1916.

31. Ozaki E, Sakimae A, Numazawa R. 1995. Nucleotide sequence of the gene for a thermostable esterase from *Pseudomonas putida* MR-2068. Bioscience, Biotechnology, and Biochemistry 59:1204–1207.

32. Yin R, Zhang X, Wang B, Jia J, Wang N, Xie C, Su P, Xiao P, Wang J, Xiao T, Yan B, Hirai H. 2022. Biotransformation of bisphenol F by white-rot fungus *Phanerochaete sordida* YK-624 under non-ligninolytic condition. Appl Microbiol Biotechnol 106:6277–6287.

33. Cunane LM, Chen Z-W, Shamala N, Mathews FS, Cronin CN, McIntire WS. 2000. Structures of the flavocytochrome *p*-cresol methylhydroxylase and its enzyme-substrate complex: gated substrate entry and proton relays support the proposed catalytic mechanism. Journal of Molecular Biology 295:357–374.

34. Hopper DJ. 1976. The hydroxylation of p-cresol and its conversion to *p*-hydroxybenzaldehyde in *Pseudomonas putida*. Biochemical and Biophysical Research Communications 69:462–468.

35. Hopper DJ, Jones MR, Causer MJ. 1985. Periplasmic location of p-cresol methylhydroxylase in *Pseudomonas putida*. FEBS Letters 182(2):485–488.

36. Rao ST, Rossmann MG. 1973. Comparison of super-secondary structures in proteins. Journal of Molecular Biology 76:241–256.

37. Londer YY. 2010. Expression of recombinant cytochromes c in *E. coli*. Heterologous gene expression in E coli: Methods and Protocols:123–150.

38. Causer MJ, Hopper DJ, McIntire WS, Singer TP. 1984. Azurin from Pseudomonas putida: an electron acceptor for p-cresol methylhydroxylase. Portland Press Ltd.

39. Alphand V, Carrea G, Wohlgemuth R, Furstoss R, Woodley JM. 2003. Towards large-scale synthetic applications of Baeyer-Villiger monooxygenases. Trends in Biotechnology 21:318–323.

40. Michelin RA, Sgarbossa P, Scarso A, Strukul G. 2010. The Baeyer–Villiger oxidation of ketones: A paradigm for the role of soft Lewis acidity in homogeneous catalysis. Coordination Chemistry Reviews 254:646–660.

41. Fürst M, Gran-Scheuch A, Aalbers F, Fraaije M. 2019. Baeyer-Villiger monooxygenases: tunable oxidative biocatalysts. ACS Catal 9: 11207–11241.

42. Torres Pazmiño DE, Baas B-J, Janssen DB, Fraaije MW. 2008. Kinetic mechanism of phenylacetone monooxygenase from *Thermobifida fusca*. Biochemistry 47:4082–4093.

43. Dudek HM, Torres Pazmiño DE, Rodríguez C, de Gonzalo G, Gotor V, Fraaije MW. 2010. Investigating the coenzyme specificity of phenylacetone monooxygenase from *Thermobifida fusca*. Applied Microbiology and Biotechnology 88:1135–1143.

44. Leipold F, Rudroff F, Mihovilovic MD, Bornscheuer UT. 2013. The steroid monooxygenase from *Rhodococcus rhodochrous*; a versatile biocatalyst. Tetrahedron: Asymmetry 24:1620–1624.

45. Mirza IA, Yachnin BJ, Wang S, Grosse S, Bergeron H, Imura A, Iwaki H, Hasegawa Y, Lau PC, Berghuis AM. 2009. Crystal structures of cyclohexanone monooxygenase reveal complex domain movements and a sliding cofactor. Journal of the American Chemical Society 131:8848–8854.

46. Chen R, Huang J, Li X, Yang C, Wu X. 2023. Functional characterization of an efficient ibuprofen-mineralizing bacterial consortium. Journal of Hazardous Materials 447:130751.

47. Ke Z, Lan M, Yang T, Jia W, Gou Z, Chen K, Jiang J. 2021. A two-component monooxygenase for continuous denitration and dechlorination of chlorinated 4-nitrophenol in *Ensifer* sp. strain 22-1. Environmental Research 198:111216.

48. Bradford MM. 1976. A rapid and sensitive method for the quantitation of microgram quantities of protein utilizing the principle of protein-dye binding. Anal Biochem 72:248–54.

49. Thomas PE, Ryan D, Levin W. 1976. An improved staining procedure for the detection of the peroxidase activity of cytochrome P-450 on sodium dodecyl sulfate polyacrylamide gels. Analytical Biochemistry 75:168–176.

50. Robert X, Gouet P. 2014. Deciphering key features in protein structures with the new ENDscript server. Nucleic Acids Res 42:W320–4.

