## Supplemental Data 1 for "Biodegradation of the endocrine-disrupting compound bisphenol F by *Sphingobium yanoikuyae* DN12"

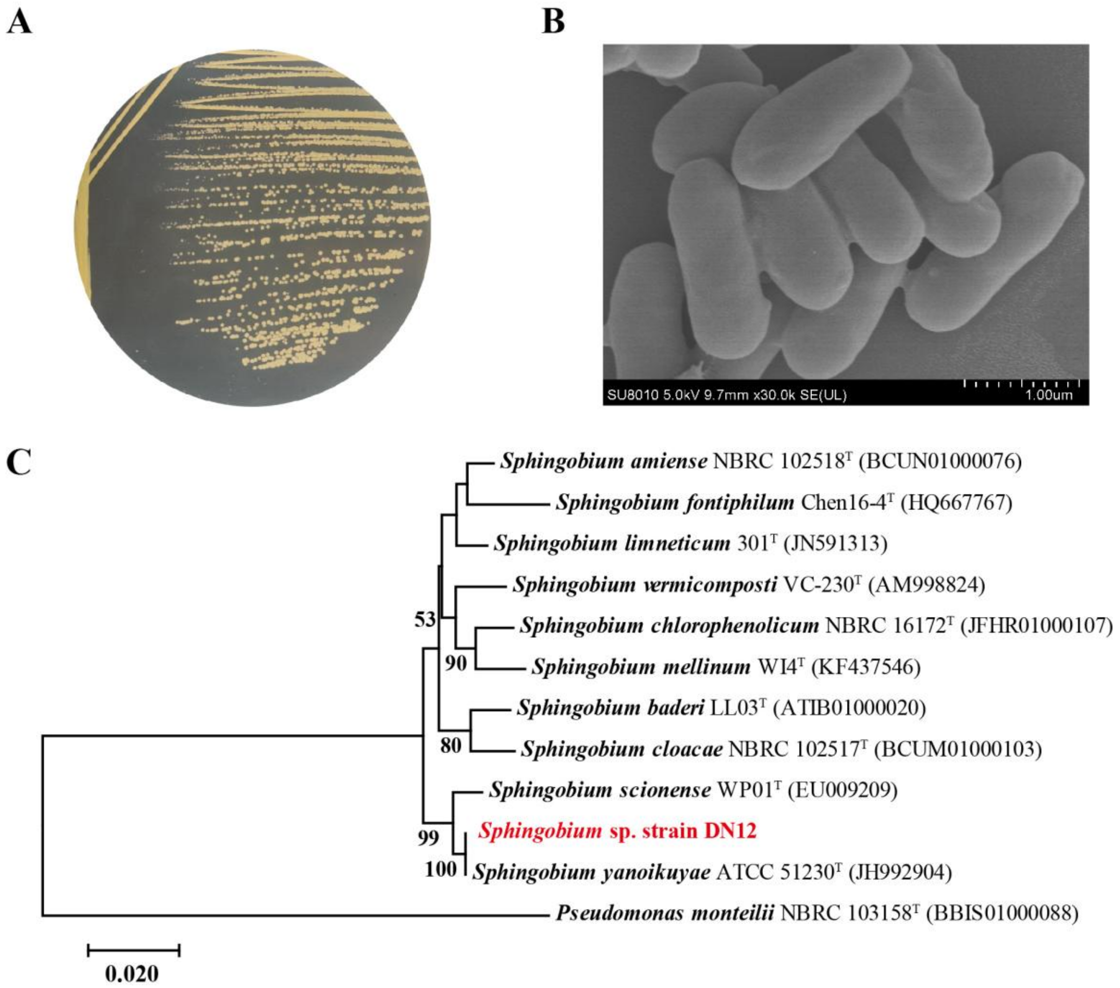


**Fig. S1** Morphology and phylogenetic analysis of strain DN12. (A) Colony morphology of strain DN12 on an LB agar plate. (B) Scanning electron micrograph of strain DN12 (scale bar = 1.0 μm). (C) Phylogenetic tree constructed based on the Neighbor-Joining method using 16S rRNA gene sequences of strain DN12 and closely related species. Bootstrap values greater than 50% (based on 1000 replicates) are shown at the branch nodes. The scale bar represents 0.020 changes per nucleotide position.

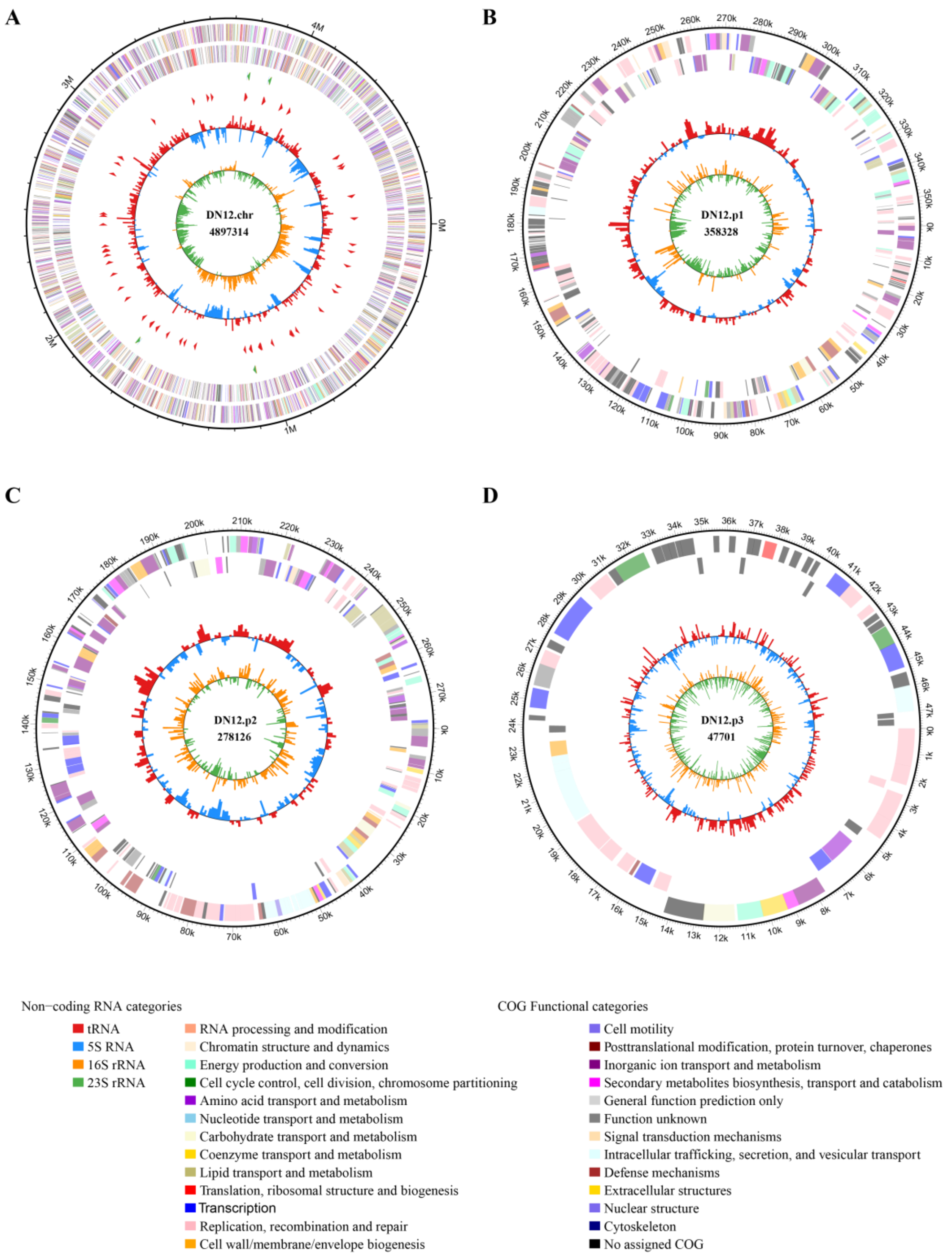


**Fig. S2** Circular representation and genetic features of the chromosome (A), pDN-1 (B), pDN-2 (C), and pDN-3 (D) of strain DN12. From outside to center: genes on the forward strand (colored by COG categories).

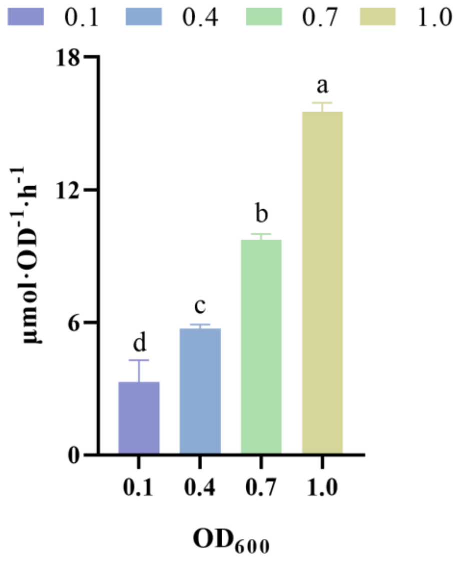


**Fig. S3** Specific degradation activities of strain DN12 at different initial inoculum densities. Specific activities were calculated as μmol BPF degraded per $OD_{600}$ per hour, based on the degradation of 0.10-0.16 mM BPF during the linear degradation phase. Bars represent the mean ± standard deviation from three biological replicates.

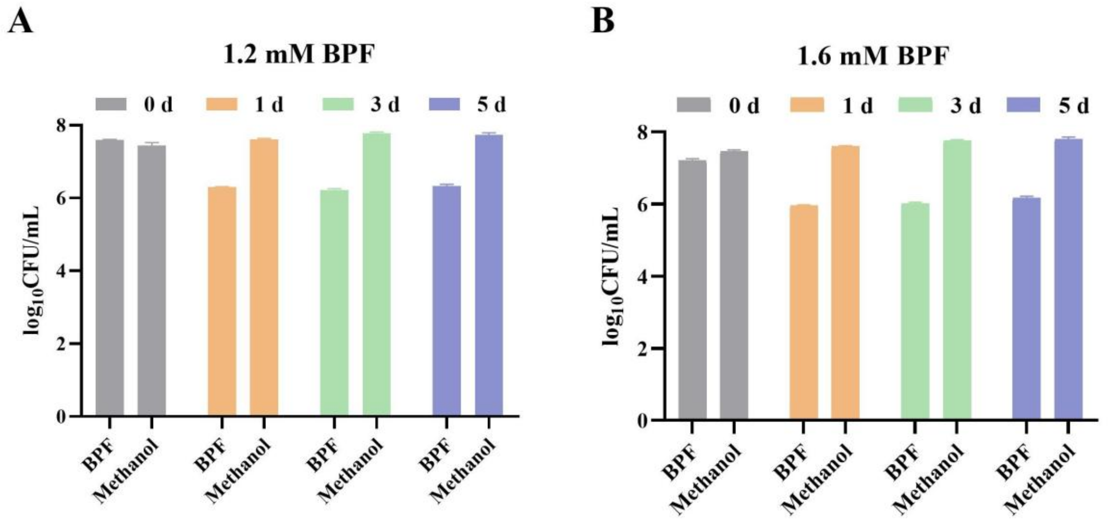


**Fig. S4** CFU-counts of strain DN12 at 0, 1, 3, and 5 days after exposure to 1.2 mM (A) and 1.6 mM (B) BPF, compared with an equal volume of methanol. Cell numbers are expressed as $\log_{10}$(CFU/mL) and represent the mean ± standard deviation of three biological replicates.

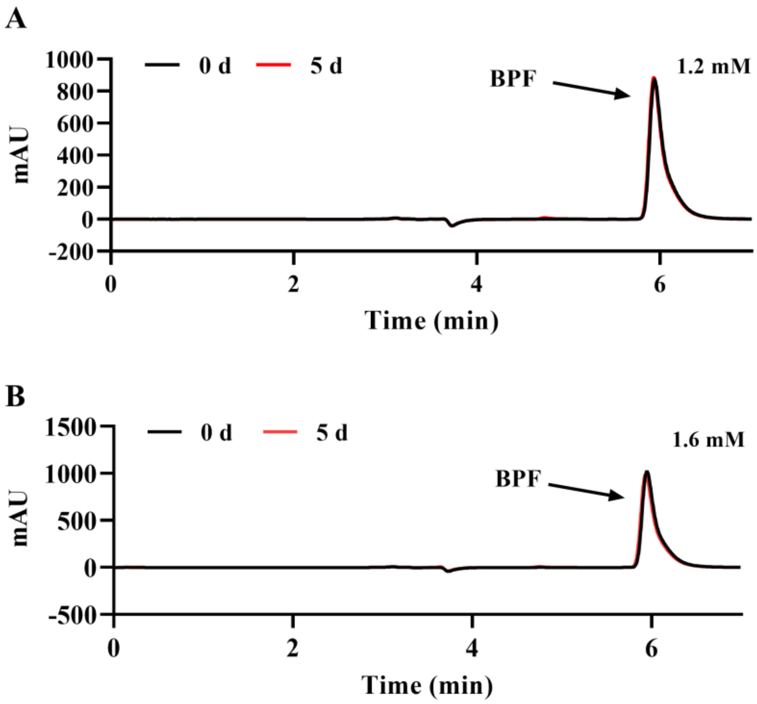


**Fig. S5** HPLC analysis of BPF degradation by strain DN12 after 0 and 5 days with 1.2 mM (A) and 1.6 mM (B) BPF.

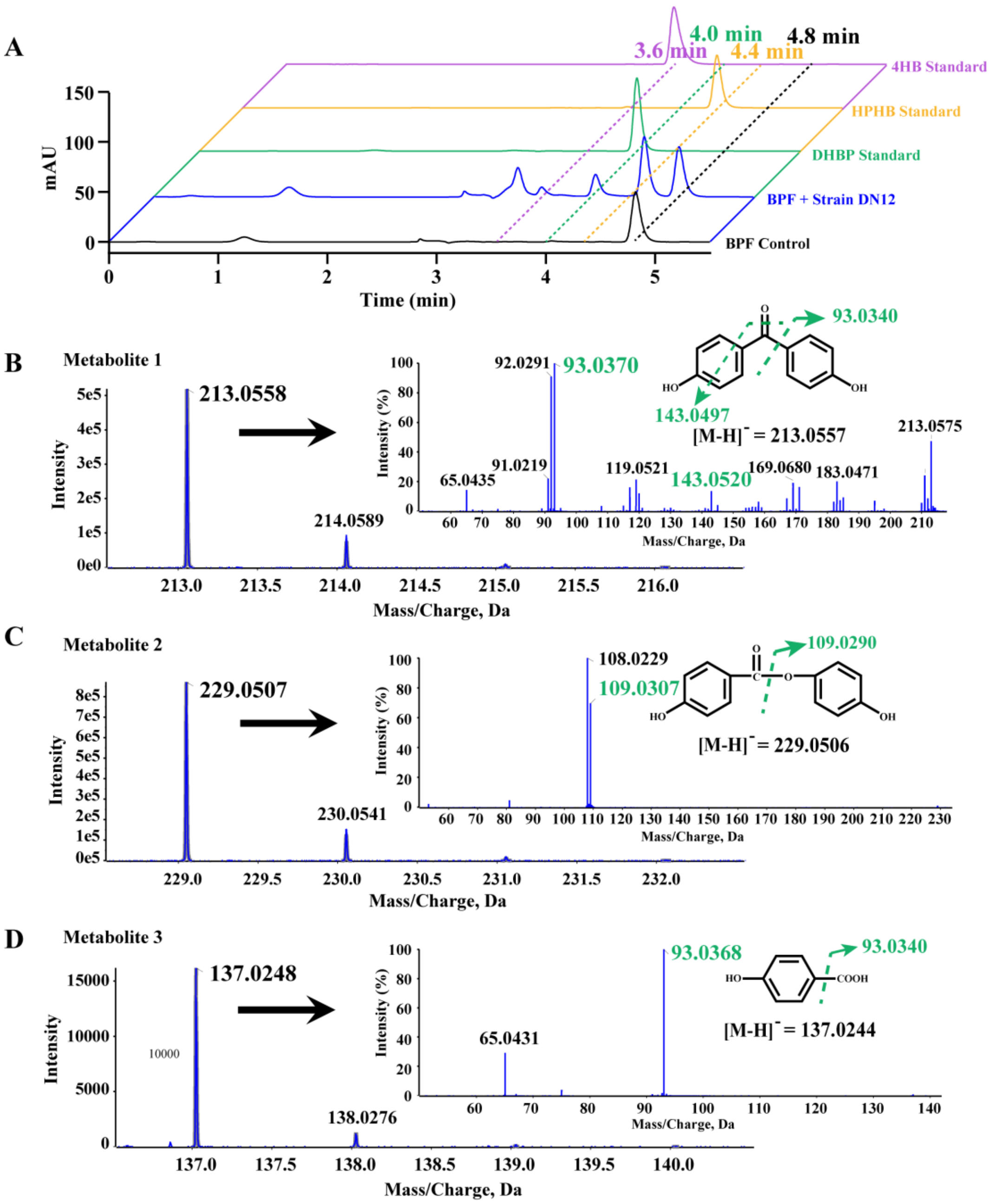


**Fig. S6** Identification of three metabolites produced during BPF degradation by strain DN12. (A) HPLC analysis of BPF degradation by strain DN12. The black line represents the control (uninoculated strain DN12), the blue line represents the BPF degradation by strain DN12, while the green, yellow, and purple lines correspond to the standards of DHBP, HPHB, and 4HB, respectively. HRMS/MS analysis of the three metabolites: DHBP (B), HPHB (C), and 4HB (D), generated during BPF degradation by strain DN12.

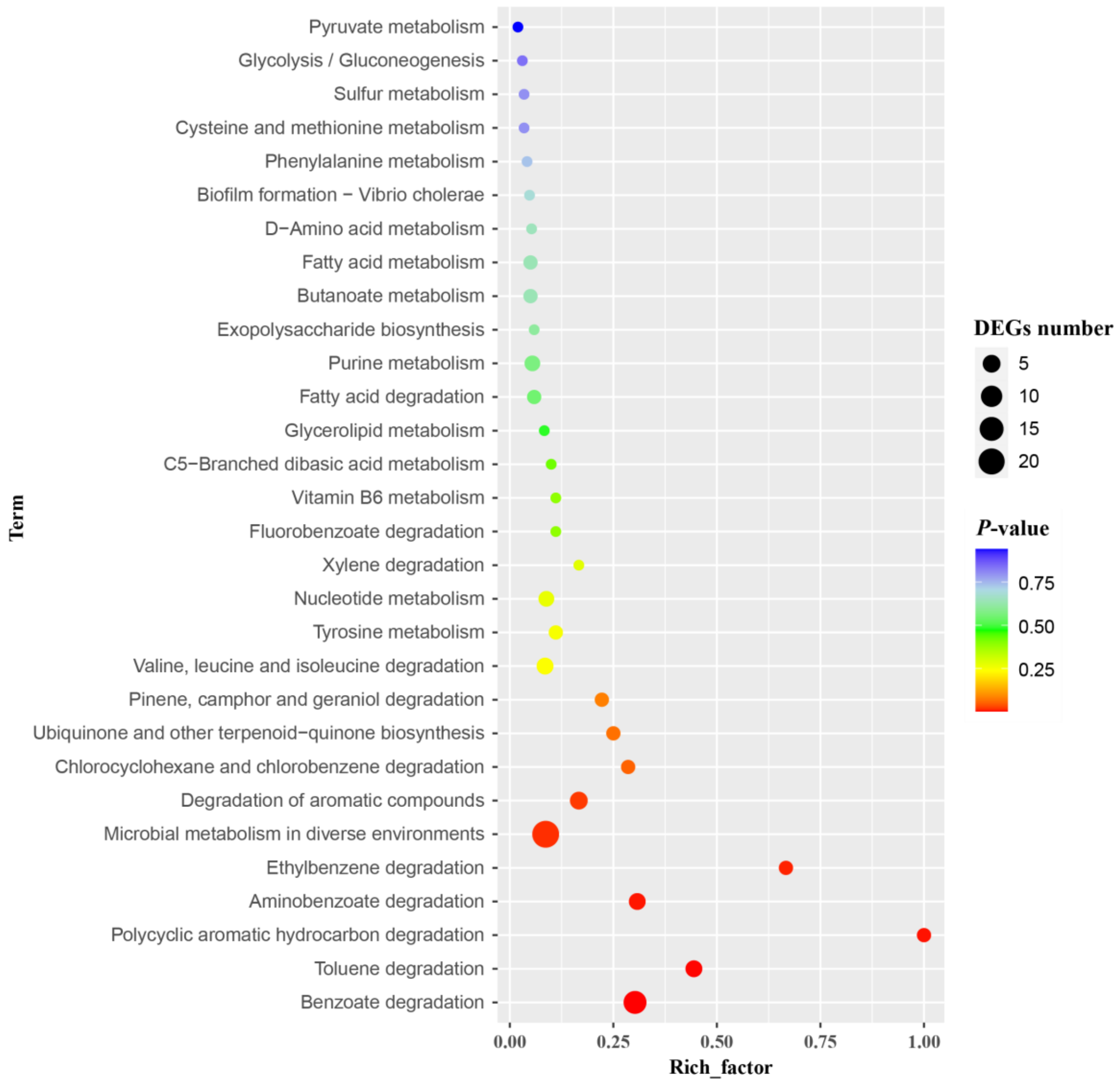


**Fig. S7** Bubble graph of the Kyoto Encyclopedia of Genes and Genomes enrichment pathway analysis. Bubble color indicates *P* value, and bubble size represents the DEG numbers.

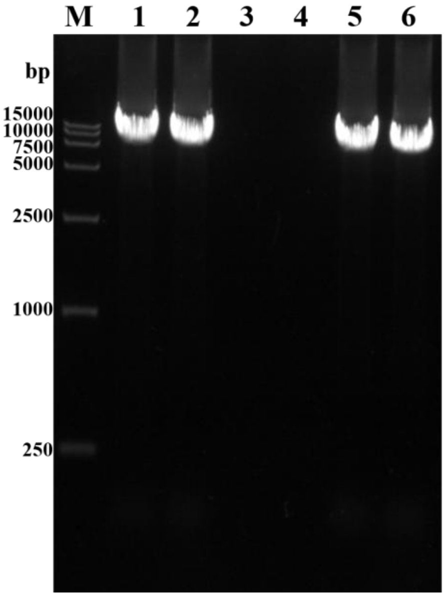


**Fig. S8** Agarose gel electrophoresis analysis of PCR products for verification of the *bpf* cluster deletion and complementation. Lane M: DL15000 DNA marker; lane 1: PCR amplification of *bpf1* from the wild-type strain DN12; lane 2: PCR amplification of *bpf2* from the wild-type strain DN12; lane 3: PCR amplification of *bpf1* from the mutant strain DN12Δ*bpf*; lane 4: PCR amplification of *bpf2* from the mutant strain DN12Δ*bpf*; lane 5: PCR amplification of *bpf1* from the complemented strain DN12Δ*bpf* (pBBR-*bpf1*); lane 6: PCR amplification of *bpf2* from the complemented strain DN12Δ*bpf* (pBBR-*bpf2*).

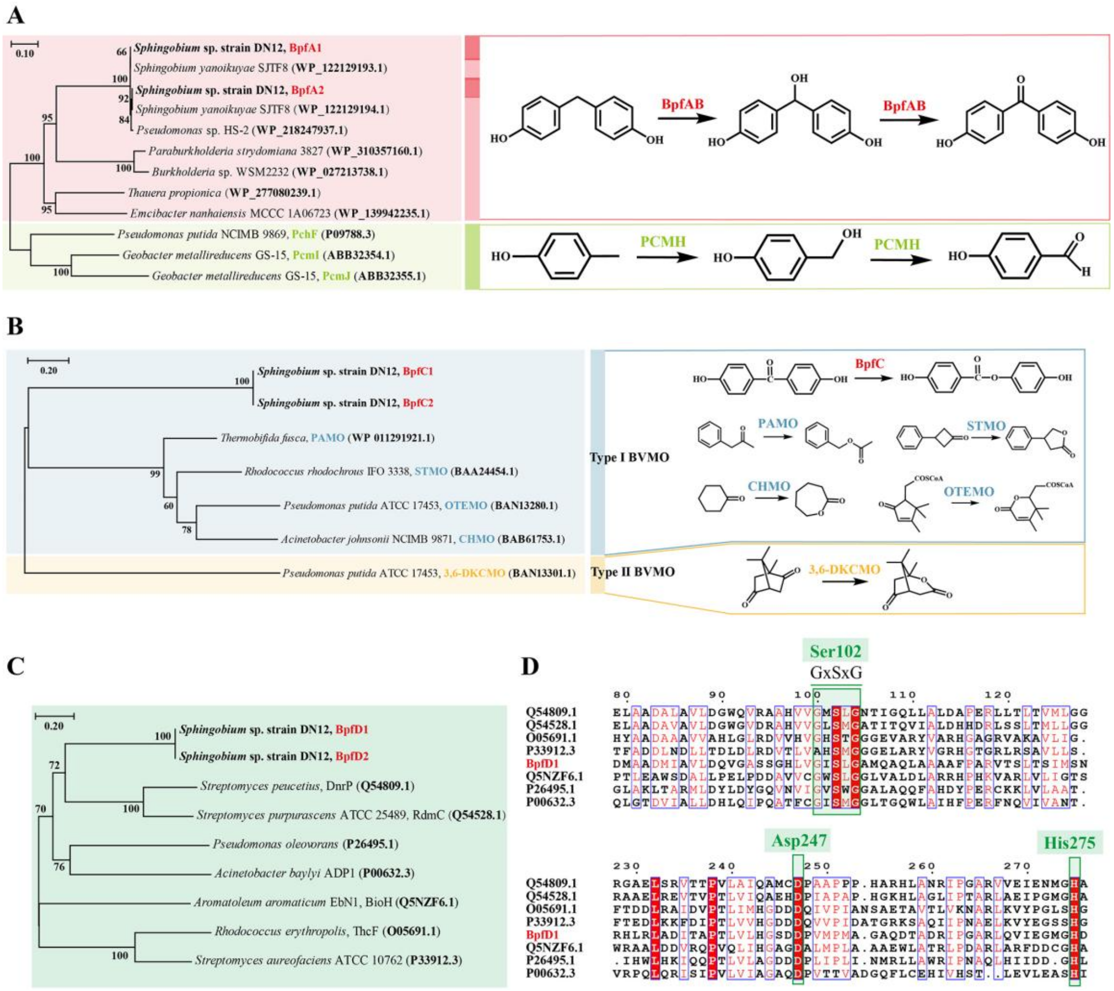


**Fig. S9** (A) Phylogenetic analysis of BpfA showing its evolutionary relationship with highly homologous proteins in the NCBI public database, as well as the types of reactions catalyzed by BpfAB and proteins with similar functions reported. (B) Phylogenetic analysis of BpfC showing its relationship to homologous proteins in the SwissProt database, including the types of reactions catalyzed by BpfC and similar enzymes. (C) Phylogenetic analysis of BpfD and its relationship to homologous proteins in the SwissProt database. (D) Amino acid sequence alignment of BpfD with closely related homologs. Conserved domains are highlighted in red, and residues proposed to contribute to BpfD substrate specificity are boxed in green.

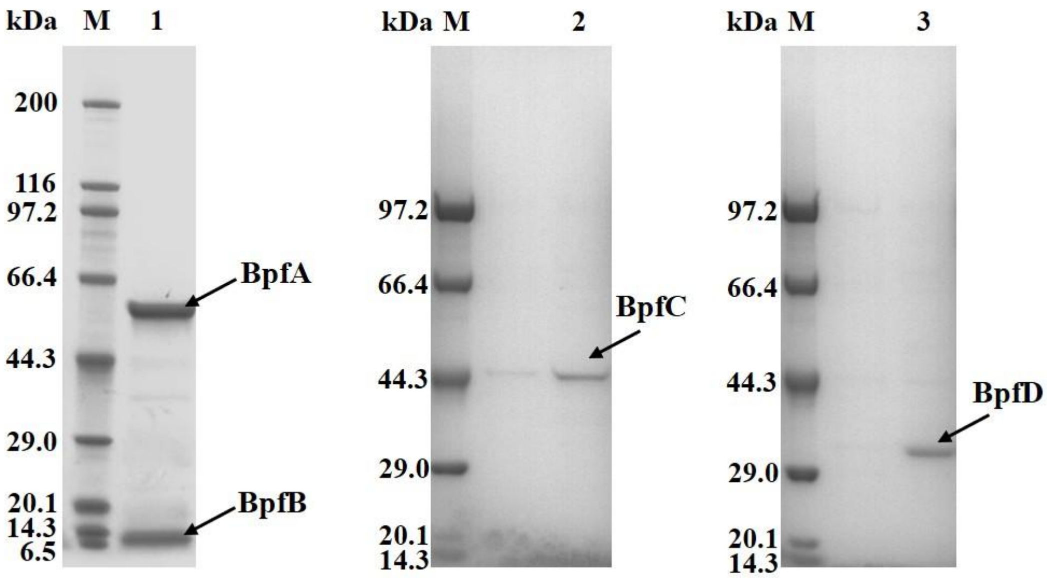


**Fig. S10** SDS-PAGE analysis of purified BpfAB, BpfC and BpfD. Lane M, molecular weight markers; lane 1, purified BpfAB; lane 2, purified BpfC; lane 3, purified BpfD.

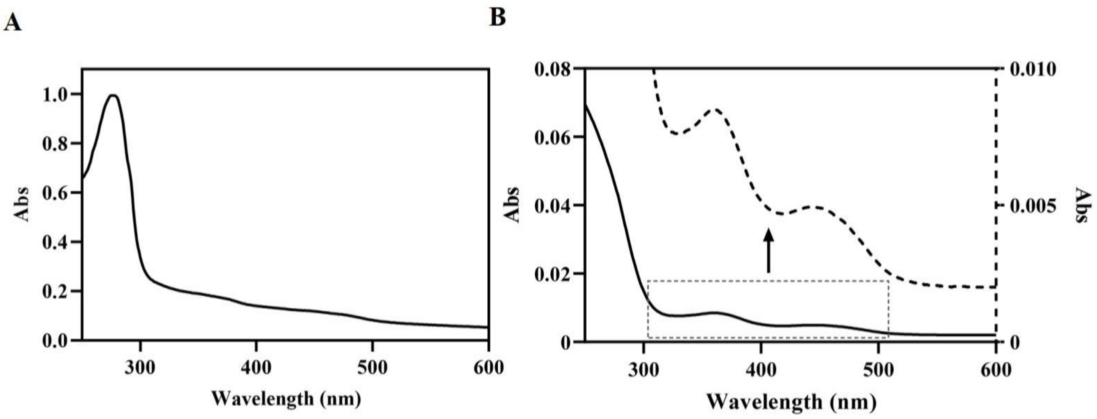


**Fig. S11** (A) UV-vis absorption spectrum of purified BpfAB. The spectrum was normalized to the absorbance at 280 nm. (B) UV-vis absorption spectrum of FAD in BpfAB, highlighting characteristic FAD absorption peaks at ~370 nm and ~450 nm.

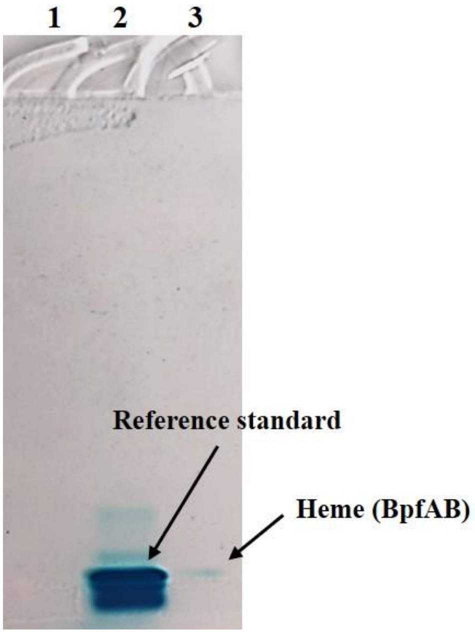


**Fig. S12** Heme staining of purified proteins using the TMB assay. Lane 1, purified BpfA (negative control); lane 2, cytochrome *c* reference standard (positive control); lane 3, purified BpfAB complex.

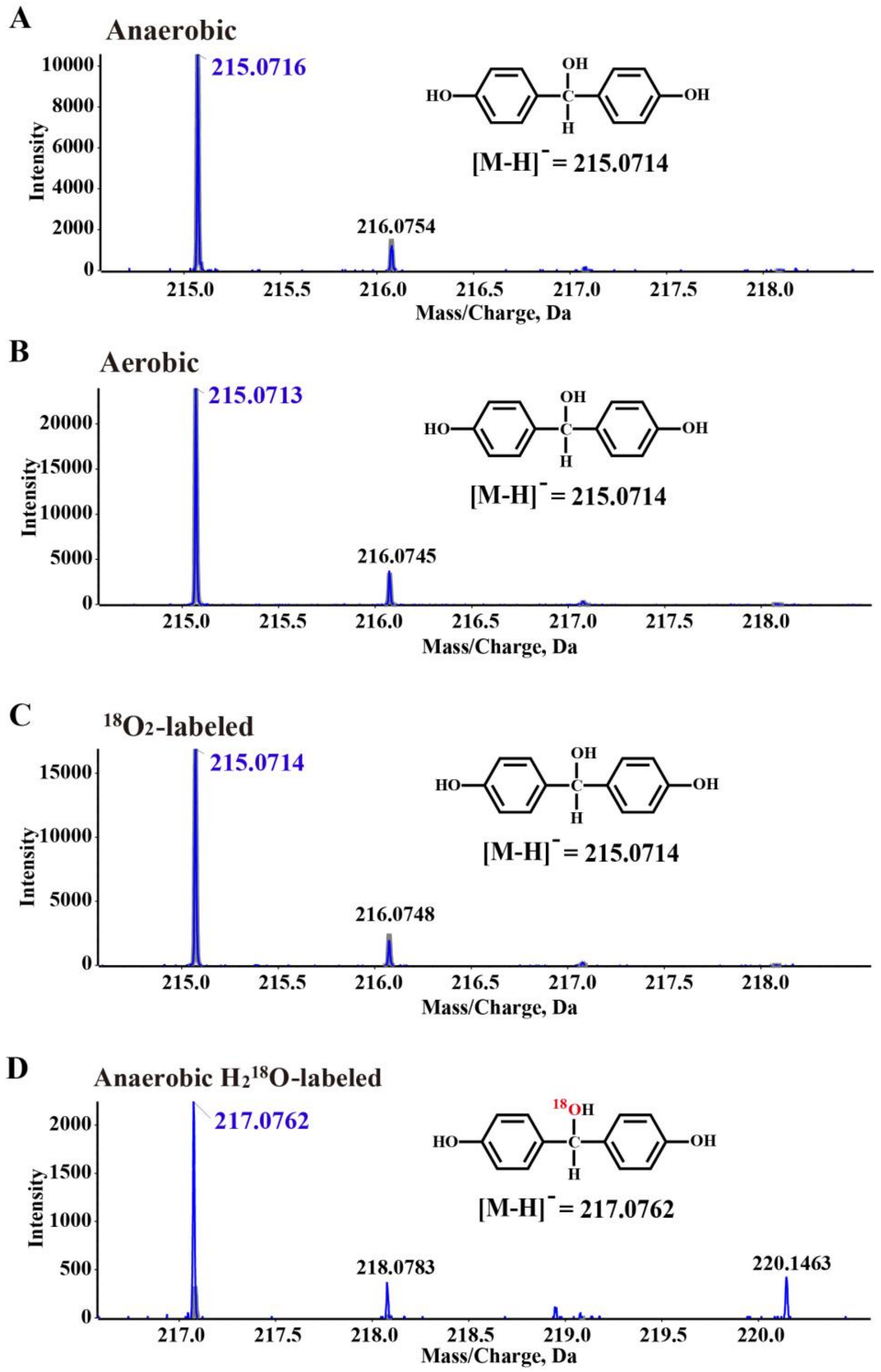


**Fig. S13.** HRMS analysis of bis(4-hydroxyphenyl)methanol, the product of BPF transformed by BpfAB under anaerobic (A), aerobic (B), $^{18}O_2$-labeled (C), and anaerobic $H_2{}^{18}O$-labeled (D) conditions.

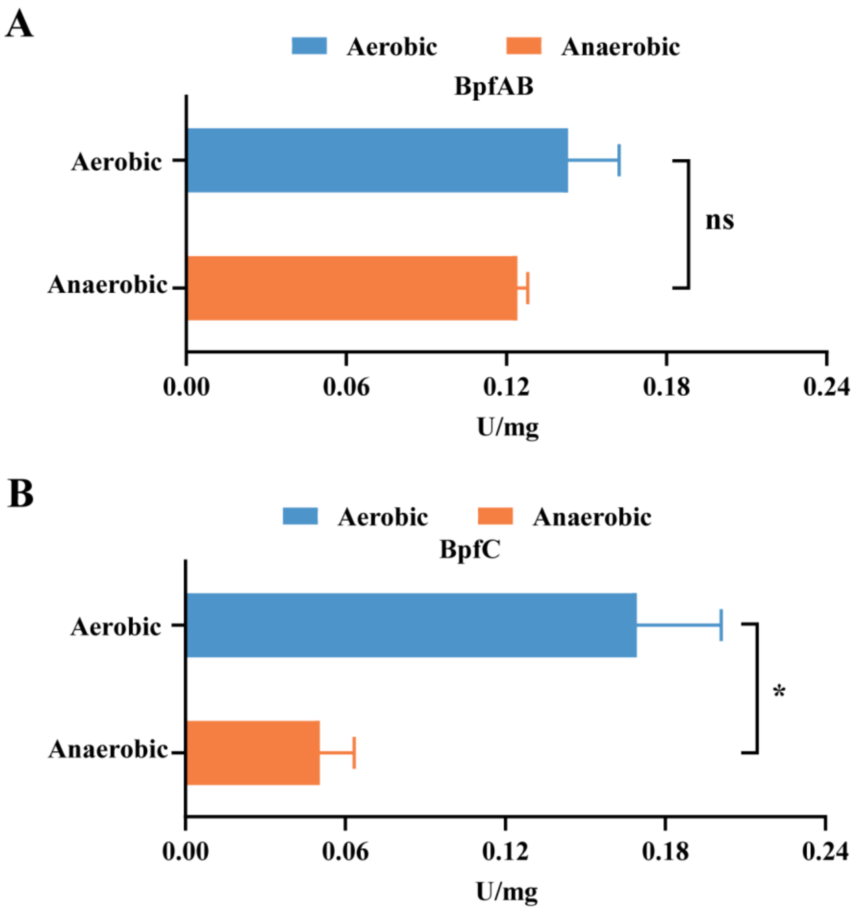


**Fig. S14** (A) Specific activity of BpfAB toward BPF under anaerobic and aerobic conditions. (B) Specific activity of BpfC toward DHBP under anaerobic and aerobic conditions. "*" indicates a significant difference (*P* < 0.05), while "ns" indicates no significant difference (*P* > 0.05). Error bars represent the standard deviation of three biological replicates.

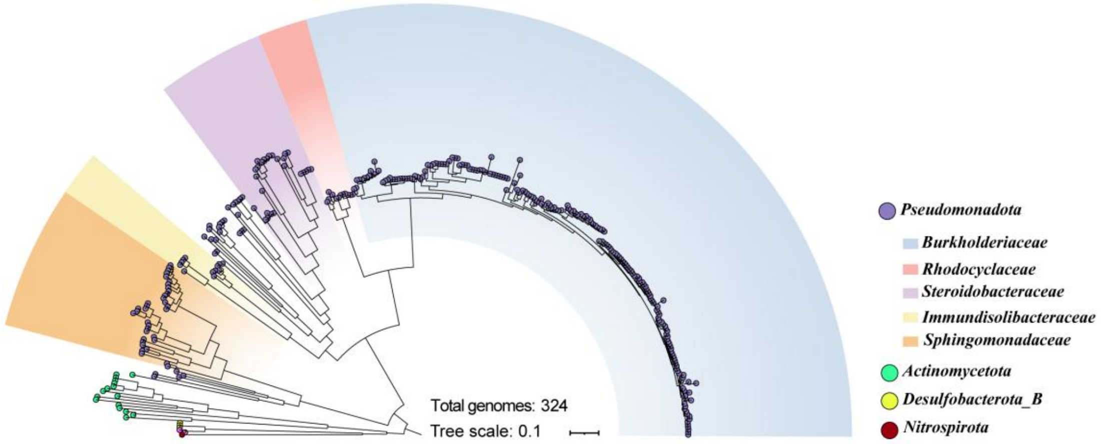


**Fig. S15** Taxonomic distribution of 324 genomes encoding BpfA-like proteins. The phylogenetic bacterial genome tree was constructed based on concatenated alignment of 120 single-copy marker genes using GTDB-Tk.

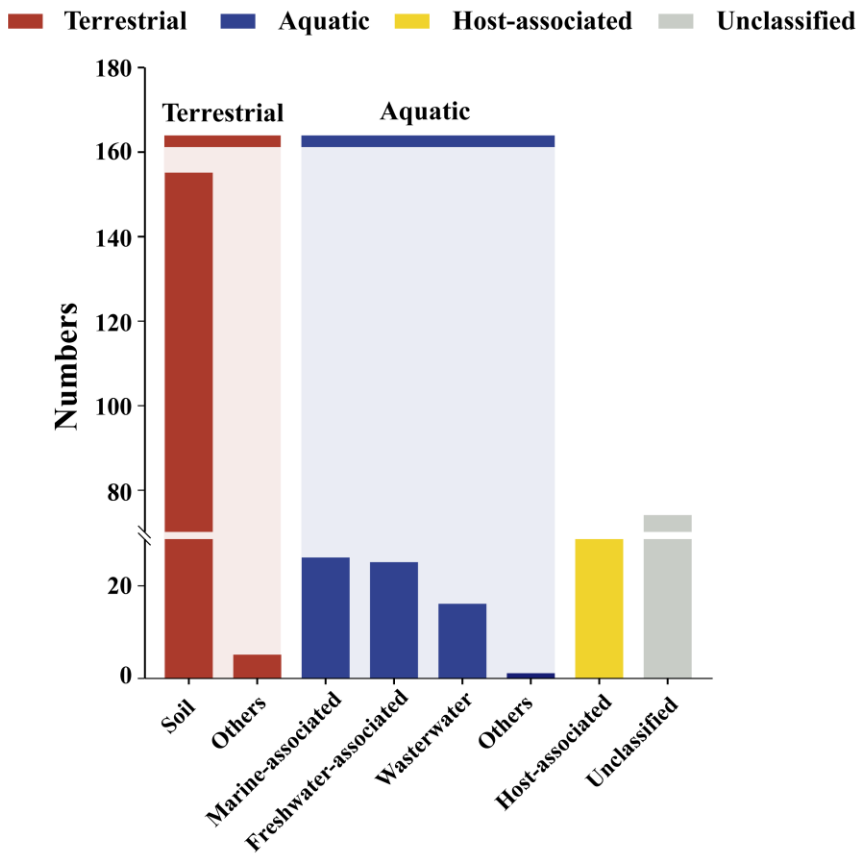


**Fig. S16** Distribution of habitat types and their corresponding numbers among 325 genomes.

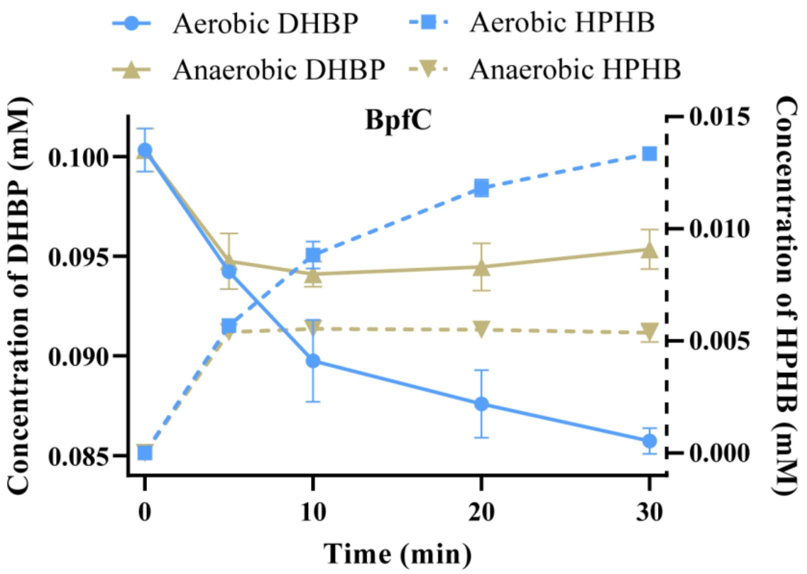


**Fig. S17** Time-course of HPHB formation from DHBP catalyzed by BpfC under aerobic and anaerobic ($N_2$-purged) conditions. Data represent the mean ± standard deviation from three independent experiments.

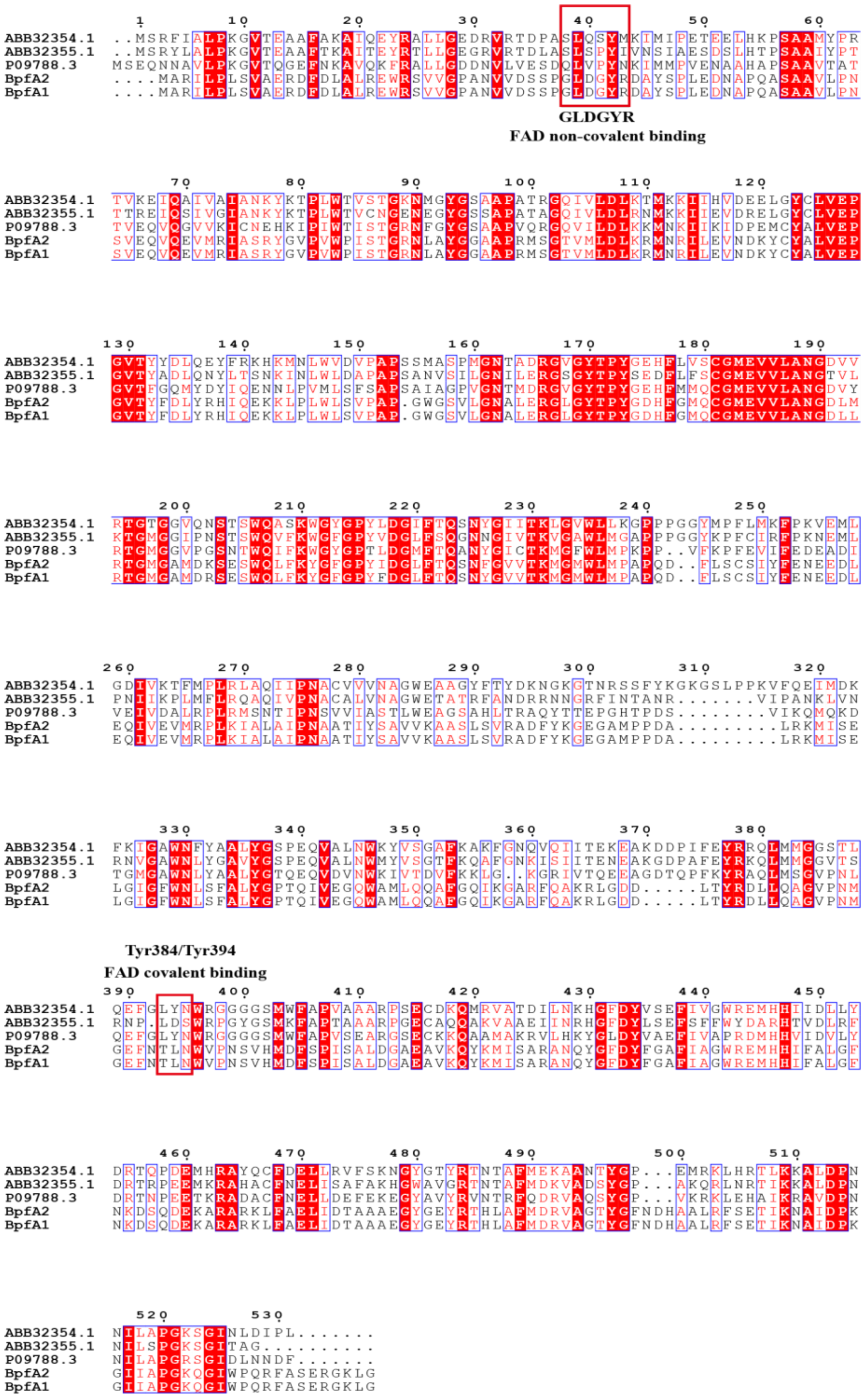


**Fig. S18** Amino acid sequence alignment of BpfA with homologous proteins PchF (*Pseudomonas putida*, P09788.3), PcmI (*Geobacter metallireducens*, ABB32354.1), and PcmJ (*Geobacter metallireducens*, ABB32355.1). Conserved residues associated with covalent FAD attachment and non-covalent FAD binding motifs are highlighted with red boxes.

**Table S1.** Sensitivity of strain DN12 to antimicrobial agents

| Antibiotics | Results[a] | Antibiotics | Results[a] |
|---|---|---|---|
| Kanamycin | - | Streptomycin | + |
| Ampicillin | - | Gentamicin | - |
| Spectinomycin | - | Tetracycline | - |
| Chloramphenicol | - | Ofloxacin | - |

[a] "-" indicates sensitivity of the strain to the antibiotic, while "+" signifies resistance.

**Table S2.** Genome information of the strain DN12.

| Features | Base number (bp) | G+C content (%) | Gene number |
|---|---|---|---|
| Genome | 5581469 | 63.23 | 5416 |
| Chromosome | 4897314 | 64.46 | 4689 |
| Plasmid 1 | 358328 | 62.53 | 381 |
| Plasmid 2 | 278126 | 62.68 | 284 |
| Plasmid 3 | 47701 | 62.11 | 62 |

**Table S3.** Degradation of bisphenol analogs by strain DN12.

| Bisphenol analogs | CAS number | Chemical structure formula | Degradation Capability [a] |
|---|---|---|---|
| BPA | 80-05-7 | 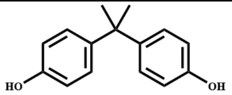 | - |
| BPS | 80-09-1 | 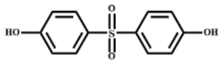 | - |
| BPF | 620-92-8 | 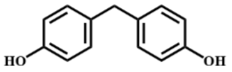 | + |
| BPB | 77-40-7 | 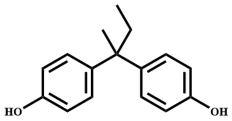 | - |
| BPE | 2081-08-5 | 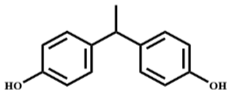 | - |
| TDP | 2664-63-3 | 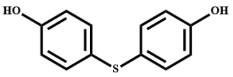 | - |
| TBBPA | 79-94-7 | 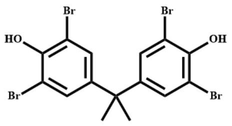 | - |
| TBBPS | 39635-79-5 | 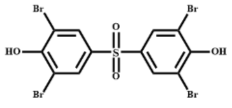 | - |

[a] The degradation efficiency of strain DN12 against various bisphenol analogs was analyzed using by HPLC after a 2-day incubation period. “+” indicates degradation; “-” signifies no degradation.

**Table S4.** Transcripts with up-regulation ($log_2$ fold change >2) in BPF-induced cells compared with those in uninduced cells

| *orf* (gene) | Product size [a] | $log_2$ fold change [b] | Database [c] | Proteins with highest identity, function, host | GenBank accession No. | Identity value |
|---|---|---|---|---|---|---|
| *orf582* | 393 | 3.95 | Swiss-Prot | *p*-hydroxybenzoate hydroxylase, PHBH, *Pseudomonas aeruginosa* PAO1 | P20586 | 60% |
| *orf583* | 256 | 2.15 | Swiss-Prot | Uncharacterized Nudix hydrolase, NudL, *Cronobacter sakazakii* ATCC BAA-894 | A7ZMT6 | 37% |
| *orf622* | 203 | 3.12 | Swiss-Prot | Epimerase family protein, YfcH, *Escherichia coli* K-12 | P77775 | 43% |
| *orf939* | 211 | 2.03 | Swiss-Prot | Aldehyde oxidoreductase iron-sulfur-binding subunit, PaoA, *Escherichia coli* O157:H7 | Q8X6I9 | 64% |
| *orf1034* | 92 | 3.33 | NR | Hypothetical protein, *Sphingobium* sp. YR657 | SHM73423 | 98% |
| *orf1658* | 92 | 3.33 | NR | Hypothetical protein, *Sphingobium* sp. YR657 | SHM73423 | 98% |

| | | | | | | |
|---|---|---|---|---|---|---|
| *orf1810* | 328 | 3.63 | Swiss-Prot | Uncharacterized oxidoreductase, YccK, *Bacillus subtilis* strain 168 | P46905 | 36% |
| *orf2927* | 313 | 3.23 | Swiss-Prot | 4-carboxy-2-hydroxymuconate-6-semialdehyde dehydrogenas, *Sphingobium* sp. SYK-6 | Q9KWL3 | 99% |
| *orf2928* | 282 | 2.79 | Swiss-Prot | Protocatechuate 4,5-dioxygenase beta chain, *Sphingobium* sp. SYK-6 | P22636 | 69% |
| *orf2929* | 134 | 2.82 | Swiss-Prot | Protocatechuate 4,5-dioxygenase alpha chain, *Sphingobium* sp. SYK-6 | P22635 | 70% |
| *orf2930* | 340 | 3.25 | Swiss-Prot | 2-keto-4-carboxy-3-hexenedioate hydratase, *Sphingobium* sp. SYK-6 | G2IQQ5 | 86% |
| *orf2931* | 298 | 2.01 | PDB | NAD-binding phosphogluconate dehydrogenase-like protein, *Alicyclobacillus acidocaldarius* subsp. *acidocaldarius* DSM 446 | 3QSG_A | 34% |
| *orf2934* | 354 | 2.51 | Swiss-Prot | (4E)-oxalomesaconate Delta-isomerase, *Novosphingobium* sp. KA1 | Q0KJL4 | 82% |

| | | | | | | |
|---|---|---|---|---|---|---|
| *orf2935* | 295 | 3.14 | Swiss-Prot | 2-pyrone-4,6-dicarboxylate hydrolase, Sphingobium sp. SYK-6 | O87170 | 87% |
| *orf3128* | 126 | 2.85 | NR | OsmC family protein, *Sphingobium yanoikuyae* | WP_125989724 | 98% |
| *orf3129* | 298 | 3.94 | Swiss-Prot | Pirin-like protein CC_3178, *Caulobacter vibrioides* CB15 | P58114 | 52% |
| *orf3130* | 106 | 3.49 | NR | Hypothetical protein, *Sphingobium yanoikuyae* | WP_374551500 | 98% |
| *orf3131* | 232 | 3.49 | Swiss-Prot | Pirin-like protein CC_1473, Caulobacter vibrioides CB15 | P58113 | 72% |
| *orf3586* | 92 | 3.33 | NR | Hypothetical protein, *Sphingobium* sp. YR657 | SHM73423 | 98% |
| *orf3687* | 337 | 3.33 | Swiss-Prot | Uncharacterized oxidoreductase, YccK, *Bacillus subtilis* strain 168 | P46905 | 36% |
| *orf5013* | 838 | 3.91 | Swiss-Prot | Vitamin B12 transporter BtuB, *Vibrio vulnificus* CMCP6 | Q8DD41 | 32% |

| | | | | | | |
|---|---|---|---|---|---|---|
| *orf5014* | 248 | 2.94 | Swiss-Prot | 3-alpha-(or 20-beta)-hydroxysteroid Dehydrogenase, *Mycobacterium tuberculosis variant bovis* AF2122/97 | P69166 | 36% |
| ***orf5015 (bpfC1)*** | 402 | 3.19 | Swiss-Prot | FAD-dependent monooxygenase, *Stachybotrys chartarum* IBT 7711 | A0A084B9Z5 | 44% |
| ***orf5016 (bpfD1)*** | 301 | 3.41 | Swiss-Prot | Aclacinomycin methylesterase, RdmC, *Streptomyces purpurascens* | Q54528 | 29% |
| *orf5017* | 189 | 2.90 | NR | Hypothetical protein, *Sphingobium yanoikuyae* | WP_122129108 | 100% |
| ***orf5018 (bpfB1)*** | 141 | 3.16 | NR | Cytochrome c, *Sphingobium yanoikuyae* | WP_122129109 | 100% |
| ***orf5019 (bpfA1)*** | 522 | 3.21 | Swiss-Prot | *P*-cresol methylhydroxylase, PCMH, *Pseudomonas putida* | P09788 | 43% |
| ***orf5022 (bpfA2)*** | 522 | 3.35 | Swiss-Prot | *P*-cresol methylhydroxylase, PCMH, *Pseudomonas putida* | P09788 | 43% |
| ***orf5023*** | 141 | 3.16 | NR | Cytochrome c, *Sphingobium yanoikuyae* | WP_122129109 | 100% |

| | | | | | | |
|---|---|---|---|---|---|---|
| ***(bpfB2)*** | | | | | | |
| *orf5024* | 189 | 2.90 | NR | Hypothetical protein, *Sphingobium yanoikuyae* | WP_122129108 | 100% |
| ***orf5025 (bpfD2)*** | 301 | 3.41 | Swiss-Prot | Aclacinomycin methylesterase, RdmC, *Streptomyces purpurascens* | Q54528 | 29% |
| ***orf5026 (bpfC2)*** | 402 | 3.15 | Swiss-Prot | FAD-dependent monooxygenase, *Stachybotrys chartarum* IBT 7711 | A0A084B9Z5 | 44% |
| *orf5027* | 248 | 2.35 | Swiss-Prot | Pyridoxal 4-dehydrogenase, *Mesorhizobium japonicum* MAFF 303099 | Q988B7 | 39% |
| *orf5031* | 496 | 2.30 | Swiss-Prot | 4-hydroxybenzaldehyde dehydrogenase (NADP(+)), *Pseudomonas putida* | Q59702 | 59% |
| *orf5044* | 377 | 2.32 | Swiss-Prot | Beta-ketoadipyl-CoA thiolase, *Pseudomonas aeruginosa* PAO1 | Q9I6R0 | 73% |
| *orf5045* | 225 | 2.74 | Swiss-Prot | Succinyl-CoA:3-ketoacid coenzyme A transferase subunit B, *Xanthomonas campestris pv. campestris str.* ATCC 33913 | P0C7I8 | 67% |

| | | | | | | |
|---|---|---|---|---|---|---|
| *orf5046* | 236 | 2.74 | Swiss-Prot | Succinyl-CoA:3-ketoacid coenzyme A transferase subunit A, *Xanthomonas campestris pv. campestris str.* B100 | B0RVK4 | 68% |
| *orf5048* | 170 | 3.76 | PDB | Hydroquinone dioxygenase small subunit, *Sphingomonas* sp. TTNP3 | 5M21_A | 100% |
| *orf5049* | 339 | 4.66 | PDB | Hydroquinone dioxygenase large subunit, *Sphingomonas* sp. TTNP3 | 5M21_B | 100% |
| *orf5050* | 328 | 4.40 | NR | hypothetical protein EBF16_01375, *Sphingobium yanoikuyae* | AYO75670 | 100% |
| *orf5051* | 498 | 4.08 | Swiss-Prot | Putative aldehyde dehydrogenase, DhaS, *Bacillus subtilis* strain 168 | O34660 | 44% |
| *orf5052* | 352 | 3.21 | Swiss-Prot | Maleylacetate reductase, *Pseudomonas* sp. P51 | P27101 | 55% |
| *orf5053* | 275 | 2.43 | Swiss-Prot | Hydroxyquinol 1,2-dioxygenase, *Rhizobium* sp. MTP-10005 | A1IIX3 | 53% |

| | | | | | | |
|---|---|---|---|---|---|---|
| *orf5055* | 193 | 4.61 | Swiss-Prot | UPF0312 protein VCM66_A0498, *Vibrio cholerae* M66-2 | C3LVF6 | 30% |
| *orf5056* | 185 | 4.64 | Swiss-Prot | Cytochrome b561 homolog 2, *Escherichia coli* K-12 | P75925 | 32% |
| *orf5057* | 203 | 4.72 | Swiss-Prot | UPF0312 protein PSPPH_0448, *Pseudomonas savastanoi pv. phaseolicola* 1448A | Q48PB8 | 36% |
| *orf5058* | 182 | 4.38 | Swiss-Prot | NAD(P)H dehydrogenase (quinone), *Azorhizobium caulinodans* ORS 571 | A8HRS7 | 83% |
| *orf5060* | 261 | 3.24 | Swiss-Prot | Pirin-like protein CC_3178, *Caulobacter vibrioides* CB15 | P58114 | 48% |

[a] Number of amino acids.

[b] Average ratio of BPF-induced to BPF-uninduced cells from three biological replicates for RNA-seq (mRNA); RPKM with $\log_2$fold change >2 (BPF-induced/uninduced) are considered significantly more abundant in the presence of BPF ($P$ <0.05).

[c] NR, NCBI Non-redundant protein sequences database; PDB, Protein Data Bank proteins.
